# Electrodiffusion dynamics in the cardiomyocyte dyad at nano-scale resolution using the Poisson-Nernst-Planck (PNP) equations

**DOI:** 10.1101/2025.03.13.642963

**Authors:** Karoline Horgmo Jæger, Aslak Tveito

## Abstract

During each heartbeat, a voltage wave propagates through the cardiac muscle, triggering action potentials in approximately two billion cardiomyocytes. This electrical activity ensures the coordinated contraction of the heart, which is essential for its pumping function. A key event in this process is the opening of voltage-gated calcium channels in the cell membrane, allowing calcium ions to enter the cardiac dyad and triggering a large-scale release of calcium ions from the sarcoplasmic reticulum through ryanodine receptors. This process is fundamental to cardiac function because calcium subsequently binds to troponin, initiating the conformational changes necessary for myofilament contraction.

The cardiac dyad is characterized by a very small volume with steep ionic concentration gradients, which is challenging for detailed mathematical modeling. Traditionally, the dyadic calcium concentration has been approximated using spatially averaged values or modeled with reaction-diffusion equations. However, at the nanometer (nm) and nanosecond (ns) scales, such approximations may be insufficient. At this resolution, the Poisson-Nernst-Planck (PNP) system provides a detailed continuous representation of the underlying electrodiffusion dynamics.

Here, we present a nano-scale computational model, representing dyad dynamics using the PNP system. Potassium, sodium, and calcium channels are incorporated in the cell membrane, along with the sodium-calcium exchanger. We demonstrate the formation of the Debye layer in the resting state and highlight how both diffusive and electrical effects are required to maintain this equilibrium. Additionally, we show that cross-species ion interactions in the dyad are electrical, and that diffusion models fail to capture this effect. Finally, we illustrate how the dyad width and diffusion coefficient influence local ionic concentrations and the timing of calcium arrival at the ryanodine receptors. These results provide new insights into the electrodiffusive properties of the dyad and clarify when solving the full PNP system is necessary for accurate modeling.

## 1 Introduction

The functioning of the heart depends on the coordinated activity of approximately two billion cardiomyocytes, [1]. Each heartbeat begins with the initiation of an action potential in the sinoatrial node, which propagates through the myocardium as a wave, depolarizing cardiomyocytes in a well-coordinated manner. At rest, cardiomyocytes maintain a transmembrane potential of approximately − 80 mV, sustained by steep ionic gradients across the cell membrane. For example, the intracellular K^+^ concentration is about 25 times higher than in the extracellular space. Since K^+^ channels remain open during the resting phase, diffusion alone would lead to a depletion of intracellular K^+^. However, electrical forces counteract this effect, maintaining the intracellular K^+^ concentration. When a depolarization wave reaches a cardiomyocyte, the transmembrane potential depolarizes, leading to the opening of voltage-gated Na^+^ channels and a rapid influx of Na^+^ ions. This further depolarizes the membrane, triggering a cascade of channel, exchanger, and transporter activations that collectively generate the full action potential.

The fundamental roles of electrical and diffusive forces across the cell membrane are well established and fully integrated into mathematical models of the action potential, [2, 3, 4]. However, in the intra- and extracellular domains, ion dynamics are often approximated using constant concentrations and voltages, see, e.g., [5], or reaction-diffusion equations, [6, 7, 8], which account for diffusion and buffering effects but neglect electrical forces. These simplifications have been very important in developing successful models of the propagating electrochemical wave in cardiac tissue, [9, 10, 11]. While such models commonly represent tissue-scale dynamics at the millimeter level, recent efforts have refined them to cellular scales, [12, 13, 14, 15, 16], and even subcellular scales down to the micrometer range, [17, 18, 19]. The integration of multiple spatial scales have also been studied, see, e.g., [20, 21, 22].

Although the Poisson-Nernst-Planck (PNP) equations provide the most accurate continuous description of electrodiffusion, they are rarely solved due to the extreme numerical resolution they require, see, e.g., [23, 24]. This raises a critical question: is it sufficient to account for electrical forces only at the cell membrane, or do these forces also influence ion dynamics in close proximity to the membrane? It is well known that near the membrane, the Debye layer forms, where ion concentrations deviate from electroneutrality and therefore cannot be accurately modeled using constant values or reaction-diffusion equations that neglect electrical effects. However, the significance of these deviations for intracellular ion dynamics remains unclear. Do these local charge imbalances merely represent a minor perturbation, or do they have tangible consequences away from the membrane? More specifically, do electrical forces within the Debye layer significantly influence the dynamics of the cardiac dyad outside the Debye layer? The purpose of this paper is to present an implementation of the PNP equations in the cardiac dyad and to analyze the electrodiffusive dynamics in this highly confined and physiologically crucial space. We solve the equations using a finite difference approach similar to [24], but here, we demonstrate how ion channels and exchangers commonly used in action potential models can be adapted to function within the PNP framework. We use the model to show how perturbations from electroneutrality decay rapidly and how the resting state is established along with its associated Debye layer. Additionally, we investigate the effects of opening individual ion channels and exchangers. For example, we investigate how the sodium-calcium exchanger (NCX) influences the dyadic dynamics when a nearby Ca^2+^ channel is open and when the Ca^2+^ channel is closed. Finally, we examine how the width of the dyad and the ionic diffusion coefficients affect the arrival time across the dyad of the Ca^2+^ wave originating from Ca^2+^ channels in the cell membrane.

## 2 Methods

### 2.1 The Poisson-Nernst-Planck (PNP) system

We apply a nano-scale computational model of electrodiffusion in the cardiomyocyte dyad based on the PNP system of equations. The system reads

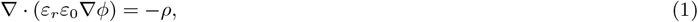

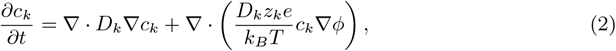

for each of the considered ion species *k*. In most of our simulations, we consider the ion species *k* = {Na^+^, K^+^, Ca^2+^, Cl^*−*^}, but in some of the first simpler examples we consider only *k* = {K^+^, Cl^*−*^}.

In the PNP system (1) and (2), *ϕ* is the electrical potential (in mV) and *c*_*k*_ is the concentration of ions of type *k* (in mM). Furthermore, *ε*_*r*_ is the relative permittivity of the medium (unitless), *ε*_0_ is the vacuum permittivity (in fF/m), *z*_*k*_ is the valence of ion species *k* (unitless), *D*_*k*_ is the diffusion coefficient of ion species *k* (in nm^2^/ms), *e* is the elementary charge (in C), *k*_*B*_ is the Boltzmann constant (in mJ/K), and *T* is the temperature (in K). Moreover, *ρ* is the charge density (in C/m^3^) defined as

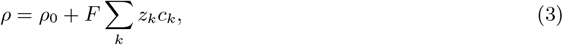

where *ρ*_0_ is the background charge density (in C/m^3^) and *F* is Faraday’s constant (in C/mol). The default values used for the model parameters in our simulations are given in Table 1.

**Table 1:**
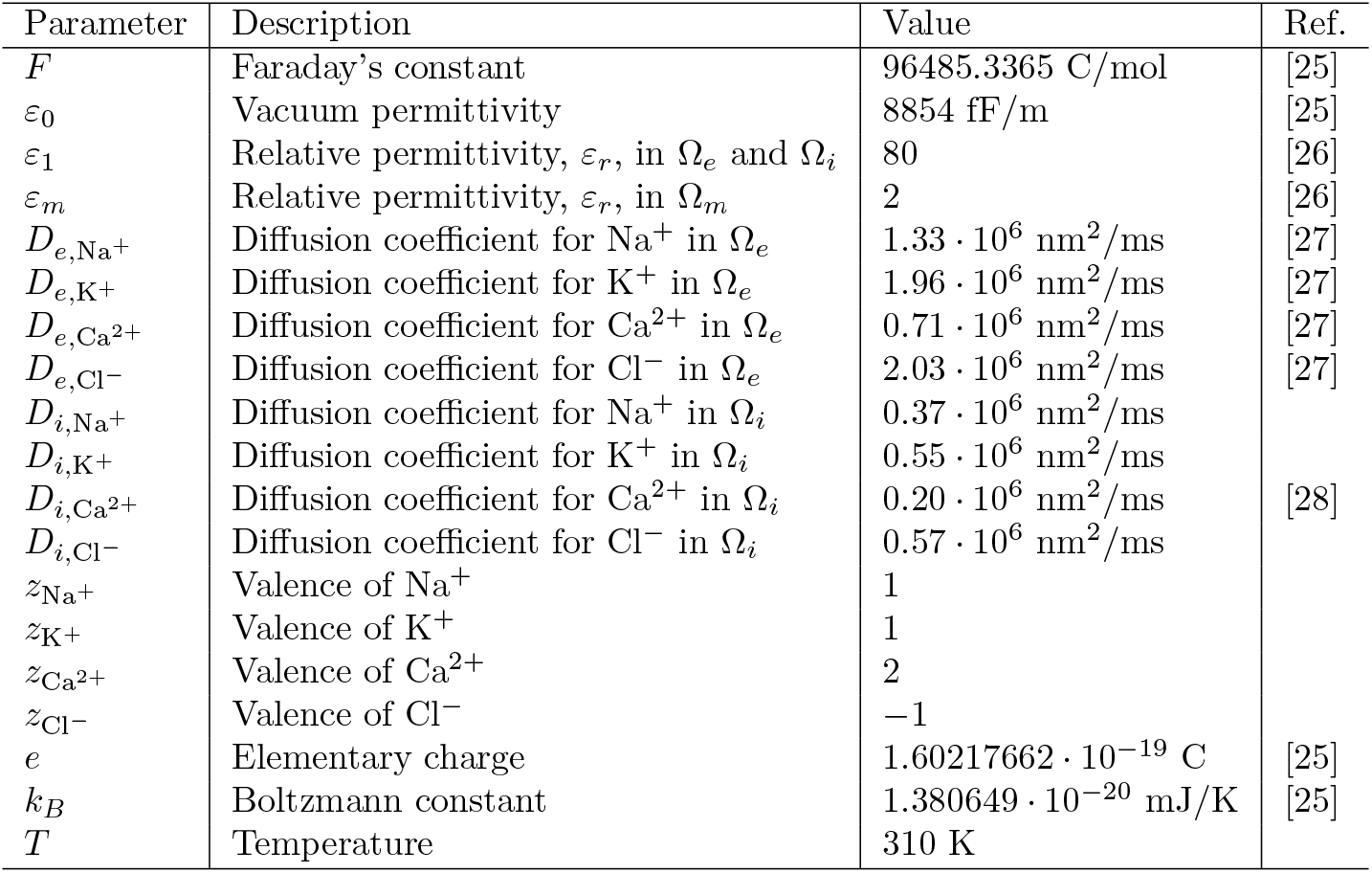
Parameter values used in the simulations. Here, Ω_*e*_ refers to the extracellular domain, Ω_*m*_ refers to the membrane domain, and Ω_*i*_ refers to the intracellular domain (see Figure 1). Note that the intracellular diffusion coefficients for Na^+^, K^+^ and Cl^*−*^ are set up such that the ratio between the intracellular and extracellular diffusion coefficients are the same as for Ca^2+^. Moreover, in the cell membrane (Ω_*m*_), the diffusion coefficient is set to zero for all ions. Electrodiffusion through channels and exchangers in the membrane is handled using local fluxes as explained in Section 2.4.

#### 2.1.1 Incorporating calcium binding buffers

As a large portion of the intracellular Ca^2+^ ions in the cardiomyocyte is bound to Ca^2+^ binding buffers, we extend the PNP model (1)–(3) to also represent two stationary Ca^2+^ binding proteins, one with high affinity and one with low affinity, based on [29]. The parameters characterizing these buffers are provided in Table 2.

**Table 2:** Parameters characterizing the intracellular Ca^2+^ binding buffers. The values are based on [29].

| Parameter | High affinity buffer | Low affinity buffer |
| --- | --- | --- |
| $B_{\text{tot}}$ | 0.20 mM | 0.56 mM |
| $k_{\text{on}}$ | 100 ms<sup>-1</sup>mM<sup>-1</sup> | 100 ms<sup>-1</sup>mM<sup>-1</sup> |
| $k_{\text{off}}$ | 0.03 ms<sup>-1</sup> | 1.3 ms<sup>-1</sup> |

The binding of an ion, *k*, to a buffer, *j*, is represented by the flux

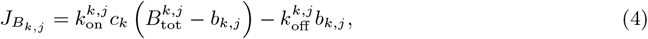

where *b*_*k,j*_ is the concentration of the ion bound to the buffer, 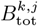 is the total buffer concentration, and 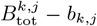 is the concentration of buffer *j* with no ion bound. Furthermore, 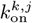 and 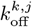 are the rate constants of the Ca^2+^ buffer binding reaction. Incorporating this reaction into the system (1)–(3), we get

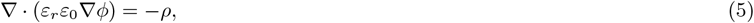

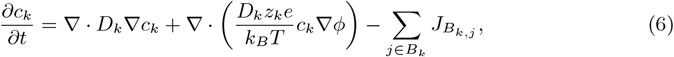

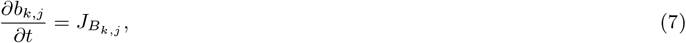

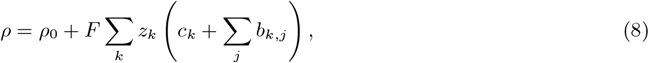

where *B*_*k*_ is a collection of all the buffers to which ion *k* may bind. In our computations, we consider two such Ca^2+^ binding buffers in the intracellular space (Ω_*i*_, see Figure 1) and no buffers for the remaining ions. Note here that in (8), *ρ*_0_ is assumed to contain the charge of the buffer proteins. We apply the model (5)–(8) in all our PNP model simulations, except for some initial simple examples using (1)–(3).

**Figure 1.**
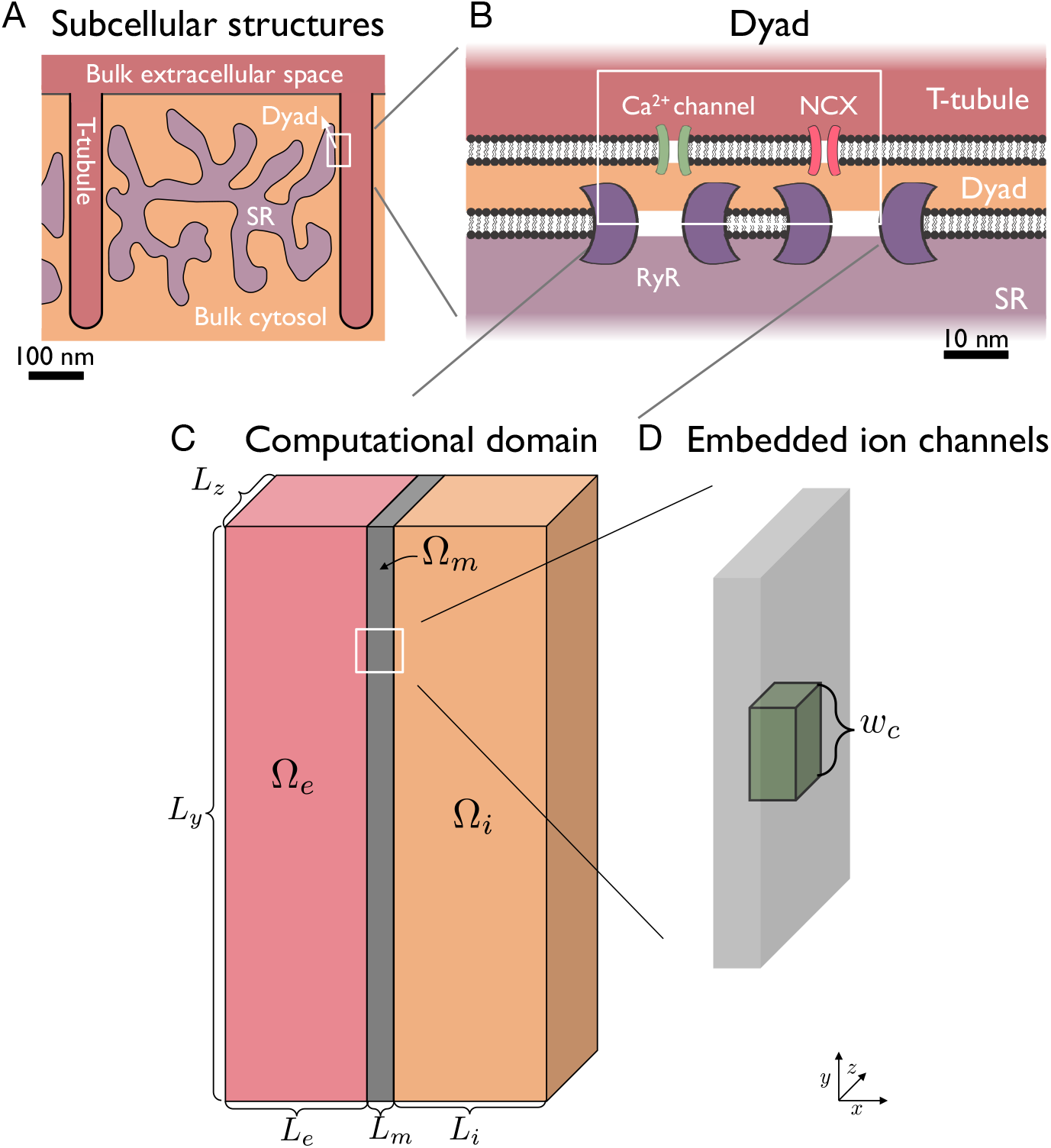
Illustration of the considered domain. A: In the cardiomyocyte, small domains called dyads are located in areas where the membrane of the SR is close to T-tubule membrane. B: In the dyad, Ca^2+^ channels and sodium-calcium exchangers (NCXs) in the cell membrane are directly apposed to ryadodine receptor channels (RyRs) in the membrane of the SR. C: The computational domain consists of an extra-cellular space, Ω_*e*_, a cell membrane, Ω_*m*_, and an intracellular domain, Ω_*i*_. A part of this domain is used to represent the dyad with an associated part of the cell membrane and extracellular space (T-tubule). D: In the computational domain, ion channels and exchangers occupy specific locations in the cell membrane.

### 2.2 Computational domain for dyad simulations

Except for the first simple examples, we consider a computational domain like the one illustrated in Figure 1C. This geometry is set up to represent:

1. The dyad (i.e., the intracellular cytosolic space between the cell membrane and the membrane of the sarcomplasmic reticulum (SR) in a location where these membranes are close and populated with Ca^2+^ channels, sodium-calcium exchangers (NCXs) and ryanodine receptors (RyRs), see Figure 1A– B)
2. The cell membrane associated with the dyad (with embedded Ca^2+^ channels and NCXs),
3. An associated part of the extracellular space (T-tubule).

The right boundary of the intracellular (dyad) part of the domain represents the SR membrane.

To represent the cell’s resting state and a realistic upstroke duration, this geometrical setup is also extended to include part of the membrane (and the associated intracellular and extracellular spaces) that do not represent the dyad location, but is rather characterized as a part of the main cell membrane. In this main cell membrane part, we include a K^+^ channel and a Na^+^ channel (see Figure 3).

**Figure 2.**
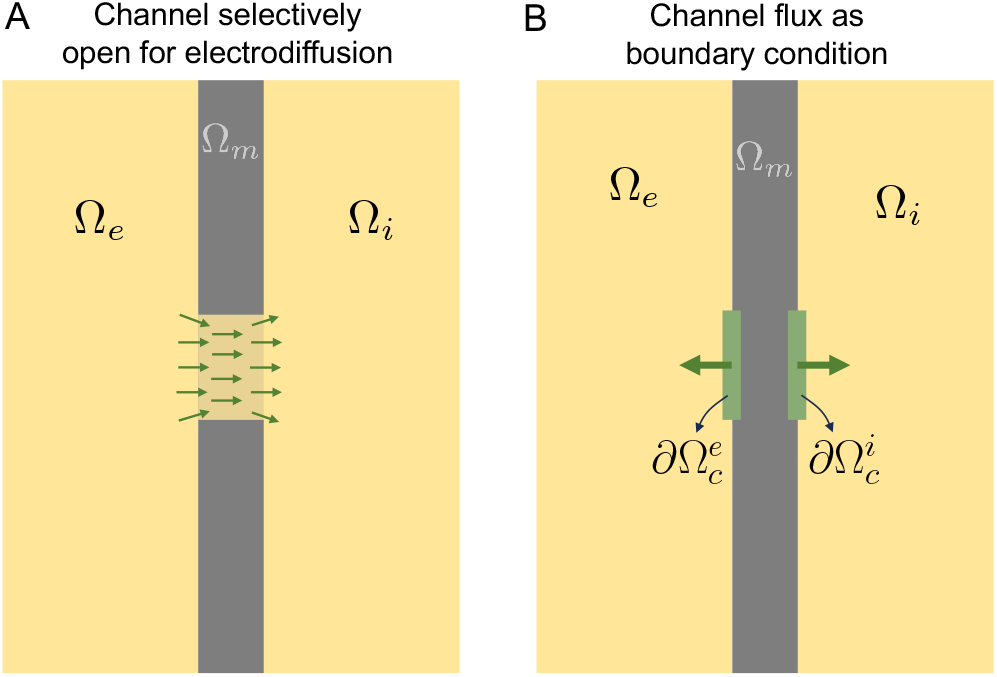
Illustration of two alternative representation of ion channels embedded in the membrane. A: Ion channels are represented as subdomains of the membrane which selectively allow for electrodiffusion of certain ion species. For example, a K^+^ channel only allows for electrodiffusion of K^+^ ions. B: The flux through ion channels are represented as internal boundary conditions for the ion concentrations associated with the channel. In this representation, electrodiffusion in the channel is formulated as a boundary condition with a specified flux. This flux can be defined by the integration of the 1D Nernst equation for the specific ion specie under consideration.

**Figure 3.**
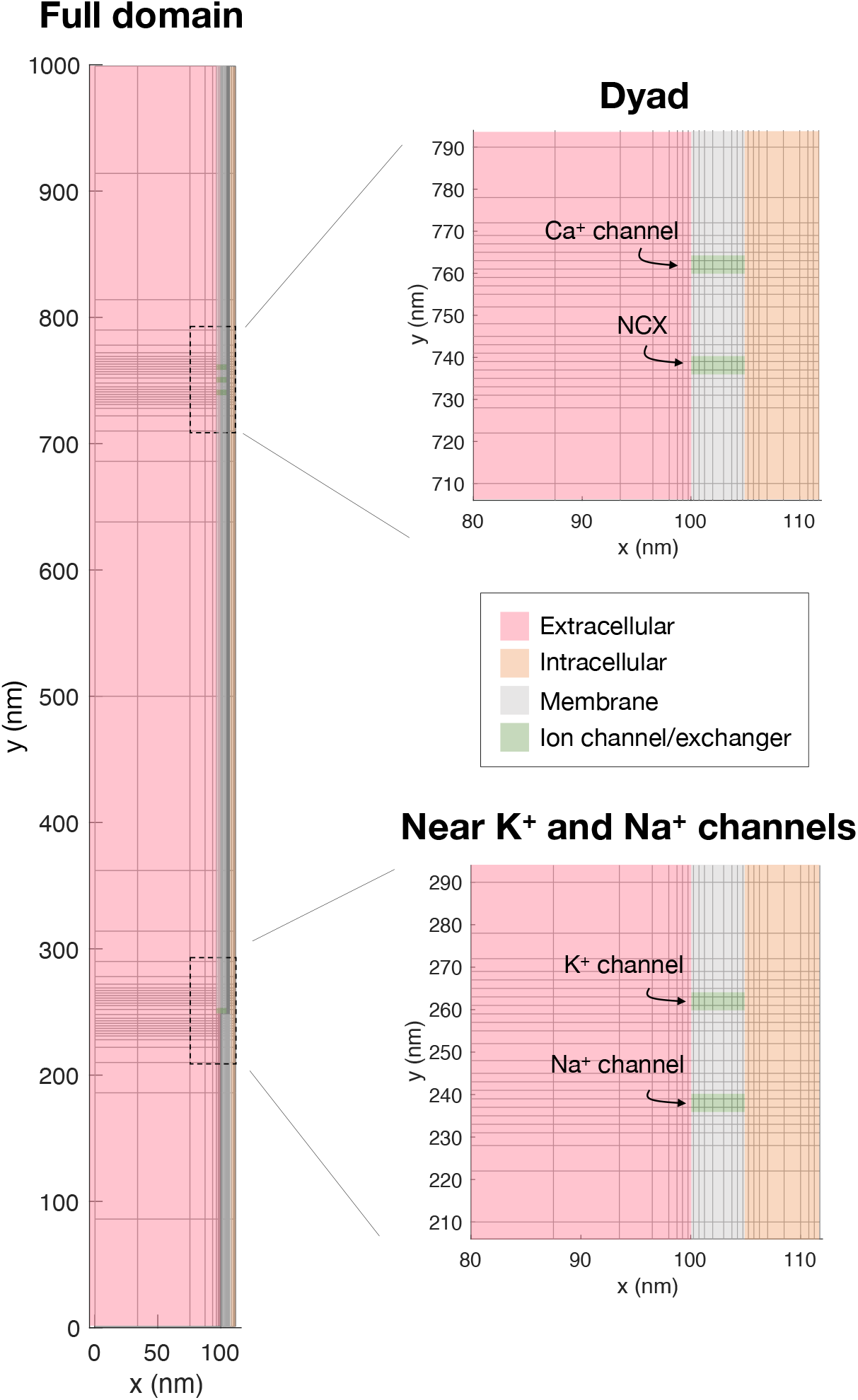
Illustration of the mesh applied in the simulations. We show a slice of the mesh in the *x*- and *y*-directions. The mesh is refined near the membrane and near membrane channels and exchangers. Similar refinements are applied in the *z*-direction. The Ca^2+^ channel and the Na^+^/Ca^2+^-exchanger (NCX) are located in a volume referred to as the dyad. A K^+^ and a Na^+^ channel are located in another volume of interest in the simulations.

The total computational domain is shaped as a rectangular cuboid, like illustrated in Figure 1C. The applied domain sizes are reported in Table 3. We let the intracellular part of the domain be denoted by Ω_*i*_, the extracellular part be denoted by Ω_*e*_, and the membrane part be denoted by Ω_*m*_.

**Table 3:**
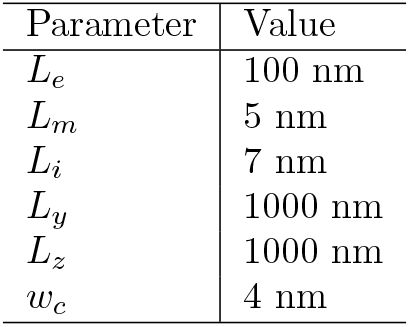
Default geometry parameter values used in the simulations. For definitions of the lengths, see Figure 1C. Note that *L*_*i*_ denotes the length from the membrane of the T-tubule to the membrane of the SR (dyad width), and will be used as a control parameter in computational experiments below.

| Parameter | Value |
| --- | --- |
| $L_e$ | 100 nm |
| $L_m$ | 5 nm |
| $L_i$ | 7 nm |
| $L_y$ | 1000 nm |
| $L_z$ | 1000 nm |
| $w_c$ | 4 nm |

As also observed in [24], the duration of the transmembrane dynamics (e.g., the duration of the upstroke) depends on the number of ion channels and the total membrane area included in the simulation. In our simulations, we consider one K^+^ channel, one Na^+^, one Ca^2+^ channel and one NCX. For this setup, we found that a membrane area of *L*_*y*_ × *L*_*z*_ = 1000 nm×1000 nm was suitable to achieve a physiologically realistic upstroke time of about 0.5 ms (see, e.g., lower right panel of Figure 9).

#### 2.2.1 Boundary conditions

For the electrical potential, *ϕ*, we apply homogeneous Neumann boundary conditions on all boundaries, except for the leftmost boundary (see Figure 1C). On that boundary we use homogeneous Dirichlet boundary conditions (*ϕ* = 0). For the ionic concentrations, we use homogeneous Neumann boundary conditions on all boundaries.

### 2.3 Initial conditions and background charge density

The initial conditions used for the ionic concentrations in the intracellular and extracellular spaces are provided in Table 4. The initial conditions for Ca^2+^ bound to the two types of intracellular Ca^2+^ binding proteins are set up such that the right-hand side of (4) is initially zero. The background charge density, *ρ*_0_, is set up such that *ρ* defined in (8) is zero for these initial conditions, hence the initial condition is always electroneutral.

**Table 4:**
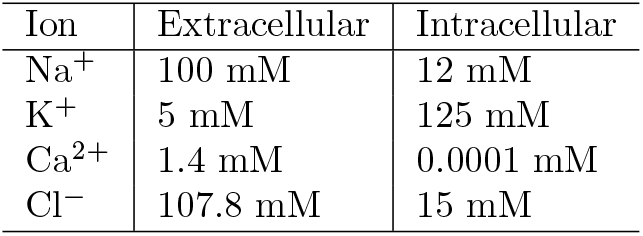
Initial conditions for the ionic concentrations in the intracellular and extracellular domains. The values are based on [27, 30, 31].

| Ion | Extracellular | Intracellular |
| --- | --- | --- |
| $\text{Na}^+$ | 100 mM | 12 mM |
| $\text{K}^+$ | 5 mM | 125 mM |
| $\text{Ca}^{2+}$ | 1.4 mM | 0.0001 mM |
| $\text{Cl}^-$ | 107.8 mM | 15 mM |

### 2.4 Representation of ion channels and exchangers

One approach for representing ion channels embedded in the cell membrane is by selectively allowing for electrodiffusion in the part of the membrane that represents the channel (see Figure 2A). For instance, for a K^+^ channel, the diffusion coefficient is zero for all ions except for K^+^ and set to some scaled parameter 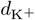 (in nm^2^/ms) for K^+^ ions. This approach for representing ion channels was applied in [24]. One disadvantage with this approach is that it might not be evident how to determine suitable diffusion coefficients or background charge densities (*ρ*_0_) for the ion channels (see [24]). Another disadvantage is that this approach is only straightforwardly implemented for ion channels, and not for membrane pumps and exchangers, like, e.g., the NCX.

To represent the fluxes through pumps and exchangers, it is more convenient to represent the fluxes as internal boundary conditions between the membrane and the intracellular and extracellular spaces (see Figure 2B). In this case, a specified expression may be directly used for the flux instead of letting the flux through the channel be governed by the PNP equations. To represent channels and exchangers in a similar manner, we will apply this alternative approach to both ion channel and exchanger fluxes in this study. However, in the Supplementary Information, we observe that the two alternative approaches illustrated in Figure 2 provide quite similar solutions (see Figures S1 and S2). The internal boundary conditions illustrated in Figure 2B take the form

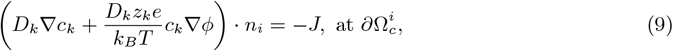

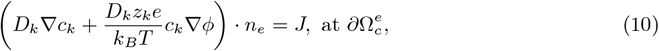

where 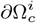 and 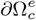 are the interfaces between the membrane and the intracellular and extracellular domains, respectively, at the location of the channel (see Figure 2B), and *n*_*i*_ and *n*_*e*_ are the outward pointing unit normal vectors of the intracellular and extracellular domains, respectively. Furthermore *J* is an expression for the channel or exchanger flux (in mMnm/ms). This flux is defined to be positive for a flux of ions in the direction from the intracellular to the extracellular space.

#### 2.4.1 Ion channel flux

The current through a single open ion channel (in yA = 10^*−*24^A) is often represented using the model

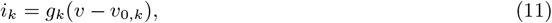

where *g*_*k*_ is the conductance of a single open channel (in zS = 10^*−*21^S), *v* is the transmembrane potential (in mV), and *v*_0,*k*_ is the Nernst equilibrium potential of the channel (in mV), see, e.g., [32]. This equilibrium potential is given by

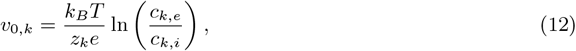

where *c*_*k,e*_ and *c*_*k,i*_ are the concentrations of ion species *k* at the extracellular and intracellular sides of the channel, respectively. This single channel current may be converted to a single channel flux (in ymol/s) by dividing the current by the ion species valence, *z*_*k*_, and Faraday’s constant, *F* :

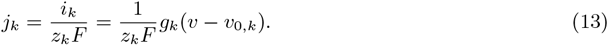

Furthermore, the single channel flux may be converted to a single channel flux density (in ymol/(nm^2^s) = mMnm/ms) by dividing the flux by the channel area (given by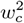, in nm^2^, see Figure 1B):

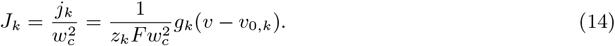

The single channel conductances, *g*_*k*_, used for the ion channels in our simulations are provided in Table 5. The transmembrane potential, *v*, is computed by

**Table 5:** Parameters for the channel and exchanger fluxes. The single channel conductances, 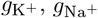, and 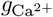 are taken from [33]. The NCX density, *δ*_NCX_, is taken from [34] and the remaining parameters characterize the NCX and are taken from [35].

| Parameter | Value | Parameter | Value |
| --- | --- | --- | --- |
| $g_{K^+}$ | 5 pS = $5 \cdot 10^9$ zS | $k_{\text{sat}}$ | 0.3 |
| $g_{\text{Na}^+}$ | 20 pS = $20 \cdot 10^9$ zS | $K_{\text{act}}$ | 0.00015 mM |
| $g_{\text{Ca}^{2+}}$ | 8 pS = $8 \cdot 10^9$ zS | $K_{\text{Ca},i}$ | 0.0036 mM |
| $\bar{I}_{\text{NCX}}$ | 4.9 A/F | $K_{\text{Ca},e}$ | 1.3 mM |
| $\delta_{\text{NCX}}$ | $4 \cdot 10^{-4}$ nm<sup>-2</sup> | $K_{\text{Na},i}$ | 12.3 mM |
| $C_m$ | 10 000 yF/nm<sup>2</sup> | $K_{\text{Na},e}$ | 87.5 mM |
| $\nu$ | 0.3 | | |

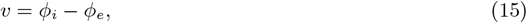

where *ϕ*_*i*_ is the average electrical potential on 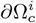, and 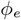 is the average electrical potential on 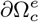 (the inlet and outlet of the channels, see Figure 2B). The concentrations *c*_*k,e*_ and *c*_*k,i*_ in the Nernst equilibrium potential (12) are similarly defined as the average ionic concentrations at 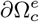 and 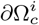, respectively.

#### 2.4.2 The sodium calcium exchanger flux

For the NCX flux, we use the formulation of the current density, *I*_NCX_, from [35], i.e.,

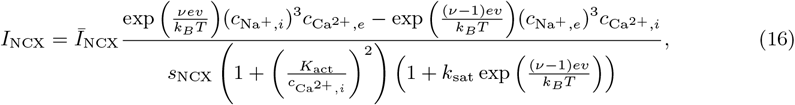

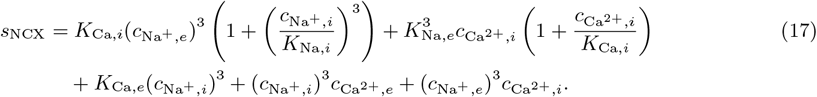

The ionic concentrations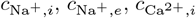, and 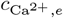 are defined as the average concentrations at 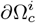 and 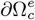 like for the ion channel fluxes. The transmembrane potential, *v*, is also defined by (15) in the same manner as for the ion channel fluxes. The parameter values involved in the channel flux are given in Table 5.

The current density, *I*_NCX_, represents the average current density over the membrane capacitance of a cardiomyocyte in units of A/F. The total current for the cell (in yA) is

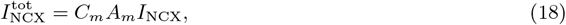

where *C*_*m*_ is the specific membrane capacitance (in yF/nm^2^) and *A*_*m*_ is the total membrane area (in nm^2^). Assuming that the membrane contains *m* NCXs, the current (in yA) through a single NCX is given by

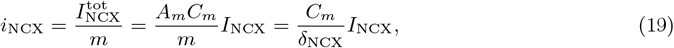

where

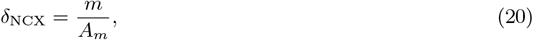

is the density of NCXs on the cell membrane.

Like for the ion channels, we may now convert this single NCX current to single NCX fluxes (in ymol/s). The current *i*_NCX_ is defined to be positive for a net positive flow of ions out of the cell. The NCX exchanges three Na^+^ ions for one Ca^2+^ ion. This means that when *i*_NCX_ is positive, Na^+^ moves out of the cell and Ca^2+^ moves into the cell, and we get

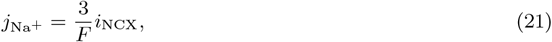

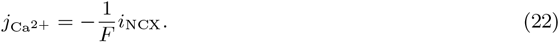

By dividing by the channel area, 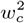, we get the flux densities (in ymol/(nm^2^s) = mMnm/ms):

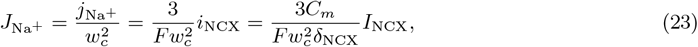

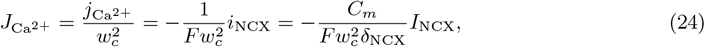

where *I*_NCX_ is defined in (16) and the parameters are found in Table 5.

### 2.5 Numerical methods

We solve the system (5)–(8) using a finite difference discretization similar to what was done in [24]. However, instead of solving (5) and (6) in two different steps, we apply a coupled scheme in this study. This allows us to increase the time step from Δ*t* = 0.02 ns used in [24] to Δ*t* = 1000 ns.

#### 2.5.1 Temporal operator splitting

We solve the system (5)–(8) using a first order temporal operator splitting technique (see, e.g., [36]) to split the buffering dynamics from the remaining system of equations. That is, for each time step, we first update *c*_*k*_ and *b*_*k*_ by solving the non-linear ordinary differential equation (ODE) system

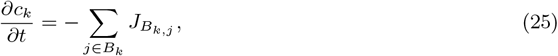

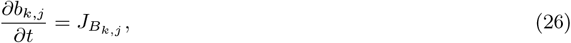

and, next, in a second step, we update *c*_*k*_ and compute *ϕ* by solving the partial differential equation (PDE) system

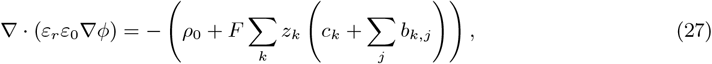

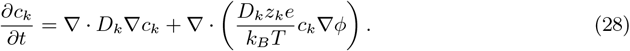

More specifically, at each time step, *n*, we assume that the solutions from the previous time step, 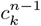 and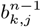, are known and use an explicit forward Euler discretization of (25) and (26) to compute 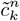 and 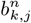.

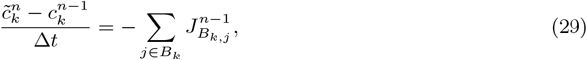

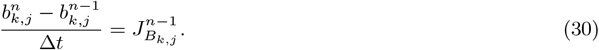

Here, 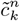 are interim solutions to be applied in the next step of the operator splitting scheme, and 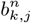 are the updated solutions of *b*_*k,j*_ for the current time step, *n*.

In the next step, we solve the system (27) and (28), in a coupled manner using an implicit backward Euler temporal discretization. To linearize the resulting system of equations, we approximate *c*_*k*_ in the term 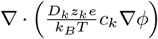 in (28) using the solution from the previous operator splitting step. This results in the following scheme for computing *ϕ*^*n*^ and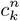 :

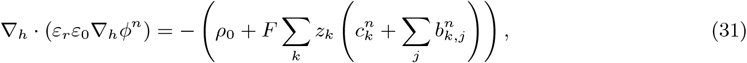

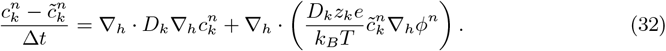

Here, ∇_*h*_ is a finite difference discretization of ∇ (see, e.g., [24]). Furthermore, 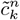 is the solution from the first step of the operator splitting scheme. If no buffers for ion *k* are present, 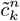 is replaced by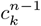.

#### 2.5.2 Discretization of membrane fluxes

For the ion channel fluxes (14), we incorporate an implicit representation of the ionic concentrations, *c*_*k*_, and the electrical potential, *ϕ*, in the second step of the operator splitting scheme. To get a linear system, we approximate the logarithm term in the Nernst equilibrium potential (12) by a Taylor series approximation around the solutions from the first operator splitting step:

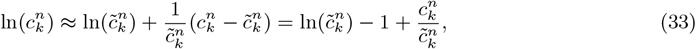

where 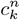 denotes the concentration of ion species *k* at time step *n*, and 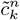 is the solution from the first operator splitting step. This yields the approximation

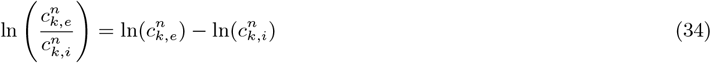

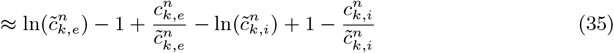

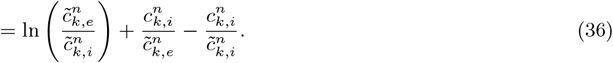

The flux (14) is thus expressed as

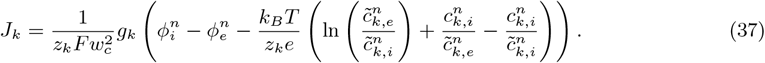

For the NCX fluxes (23) and (24), we use an explicit representation of the potential and ionic concentrations. In other words, we apply the concentrations from the previous operator splitting step,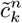, and the electrical potential from the previous time step, *ϕ*^*n−*1^.

#### 2.5.3 Mesh

To reduce the computational costs, we use a simple adaptive meshing approach, like in [24]. In this approach, a high resolution is applied near the membrane and the ion channels and a coarser resolution is applied elsewhere. A slice of the mesh in the *x*- and *y*-directions are illustrated in Figure 3. Near the membrane and the right boundary of the domain (representing the SR membrane), the distance between grid points is 0.5 nm in the *x*-direction, and the distance doubles for each grid point further away from the membrane. Similarly, the distance between grid points is 2 nm in the *y*- and *z*-directions near the channels or exchangers and doubles for each grid point as the distance from the channels or exchangers increases.

### 2.6 Visualization of simulation results

In this section, we describe the setup used to visualize the results of our simulations. To this end, Figure 4 illustrates the general setup used in the simulation visualizations.

**Figure 4.**
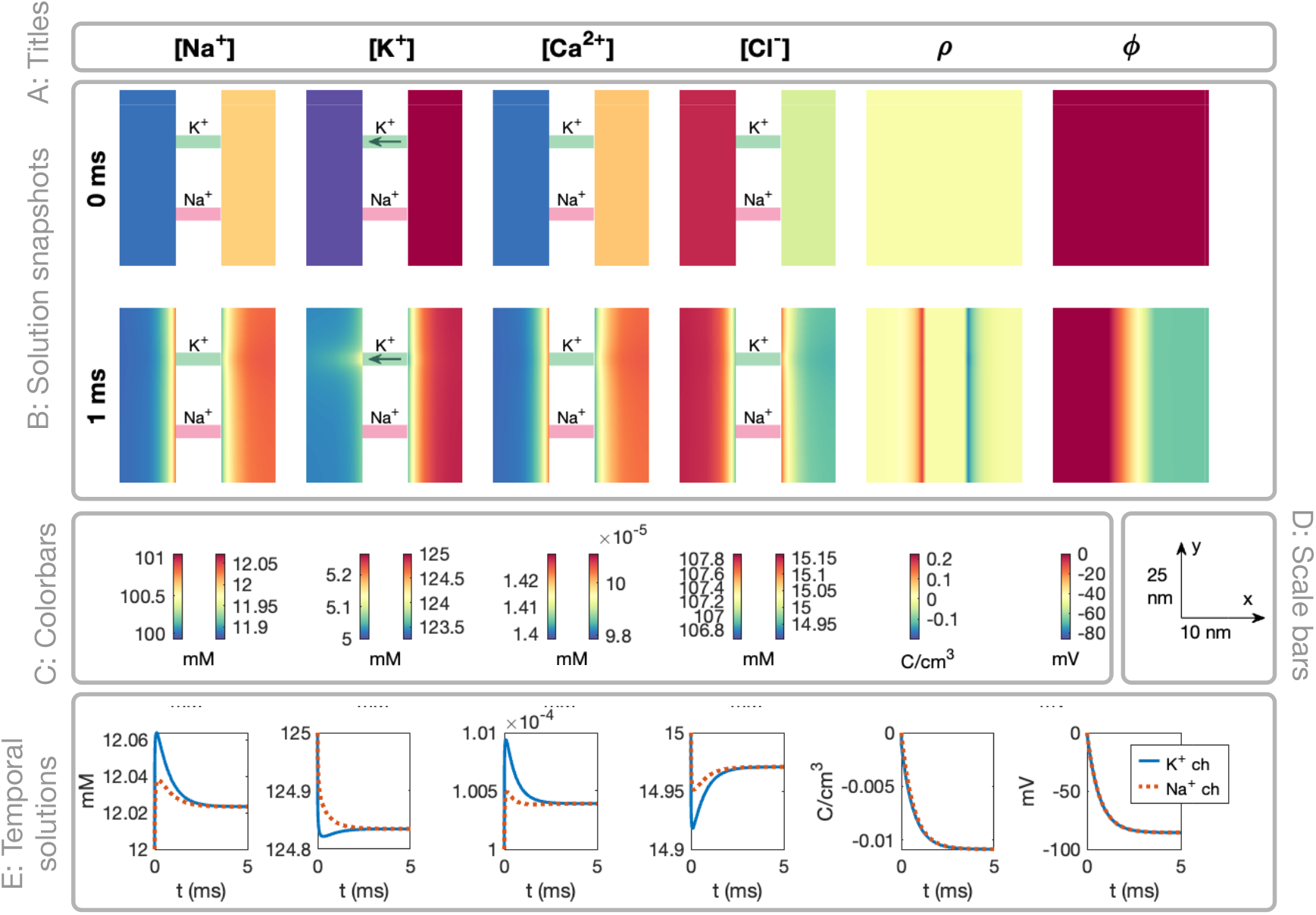
Example illustration of the setup used to visualize the simulation results. The setup is described in detail in Section 2.6, and re-used in many figures below.

#### A: Titles

In the upper (title) panel of the figure, each of the considered ionic species are listed, as well as *ρ* and *ϕ*. All figure panels below these titles depict the concentration of the ion species, or *ρ* or *ϕ*, named in these upper titles.

#### B: Solution snapshots

In the next two plot rows, snapshots of the solutions are displayed. The plot titles (Panel A) describes the variable displayed in each column. For each plot row, one point in time is considered. This time point is reported on the left side of each row. In the example illustration in Figure 4, only two time points are included (*t* = 0 ms and *t* = 1 ms), but for the actual result figures five time points are generally included.

The snapshots show the solutions in a sheet in the *x*-*y*-plane in the center of the domain in the *z*-direction. Moreover, we only consider a part of the domain that is in close proximity to channels of interest. Figure 4 shows the dynamics close to the K^+^ and Na^+^ channels, but some of the result figures focus on the dyad area (close to the Ca^2+^ channel and NCX) instead.

In the plots of the ionic concentrations, the part of the domain representing the membrane is removed from the visualizations (because all concentrations are zero in the membrane). Instead, illustrations of the ion channels and exchangers are included. In these illustrations, a green color indicates that the channel is open and a red/pink color indicates that the channel is closed. Furthermore, for the ionic species that is able to move through the channel or exchanger, an arrow indicating the direction of flow of the ions is also included. The extracellular space is illustrated on the left side of the channels and the intracellular space is illustrated on the right side of the channels. For *ρ* and *ϕ* the membrane is included in the solution snapshots. Thus, we do not explicitly display the ion channels and exchangers in these plots, but they are located at the locations indicated in the concentrations plots.

#### C: Colorbars

Below the solution snapshots, colorbars for these solutions are included. For *ϕ* and *ρ* the same colormap scaling is used in the extracellular, intracellular, and membrane domains, and thus only one colorbar is included for each of these two variables. For the ionic concentrations, on the other hand, the colormap scaling is different in the intracellular and extracellular domains. Therefore, two colorbars are provided for each ionic species. The right colorbar shows the colormap scaling in the extracellular space and the left colorbar shows the colormap scaling in the intracellular space. The colorbar units are provided below the colorbars.

#### D: Scale bars

On the right-hand side of the colormaps, scalebars for the snapshots are provided for the *x*- and *y*-directions.

#### E: Temporal solutions

In the bottom row of the figures, the temporal evolution of the solutions in two spatial points are plotted. More specifically, we consider the points located in the center of the intracellular domain in the *x*-direction and at the location of two considered channels in the *y*-and *z*- directions. For the solutions displayed in Figure 4, this corresponds to the solutions 3.5 nm to the right of the K^+^ and Na^+^ channels.

## 3 Results

In this section, we describe the results of simulations using the model described in the Methods section. First, we will demonstrate properties of the PNP model using two simplified examples. Next, we will show solutions of the PNP model set up to represent the dyad. For instance, we show results of simulations including the opening of different types of channels and investigate how altering the diffusion coefficient or the dyad width affect the results. Finally, we will compare the PNP model solutions to the solution of alternative models for the Ca^2+^ dynamics in the dyad.

### 3.1 PNP model solutions of decay following perturbations from electroneutrality

To illustrate properties of the PNP model, we first consider a simple example of a perturbation from electroneutrality. We consider the PNP model (1)–(3) in two dimensions (2D) for the two ions K^+^ and Cl^*−*^. We let *ρ*_0_ = 0 in the entire domain. At the boundary of the domain, we apply Dirichlet boundary conditions fixing the potential, *ϕ*, at 0 mV and both ionic concentrations at 100 mM. The initial conditions are illustrated in Figure 5A. For these initial conditions, the concentrations do not initially fulfill electroneutrality (i.e., we have *ρ ≠* 0, where *ρ* is defined in (3)).

**Figure 5.**
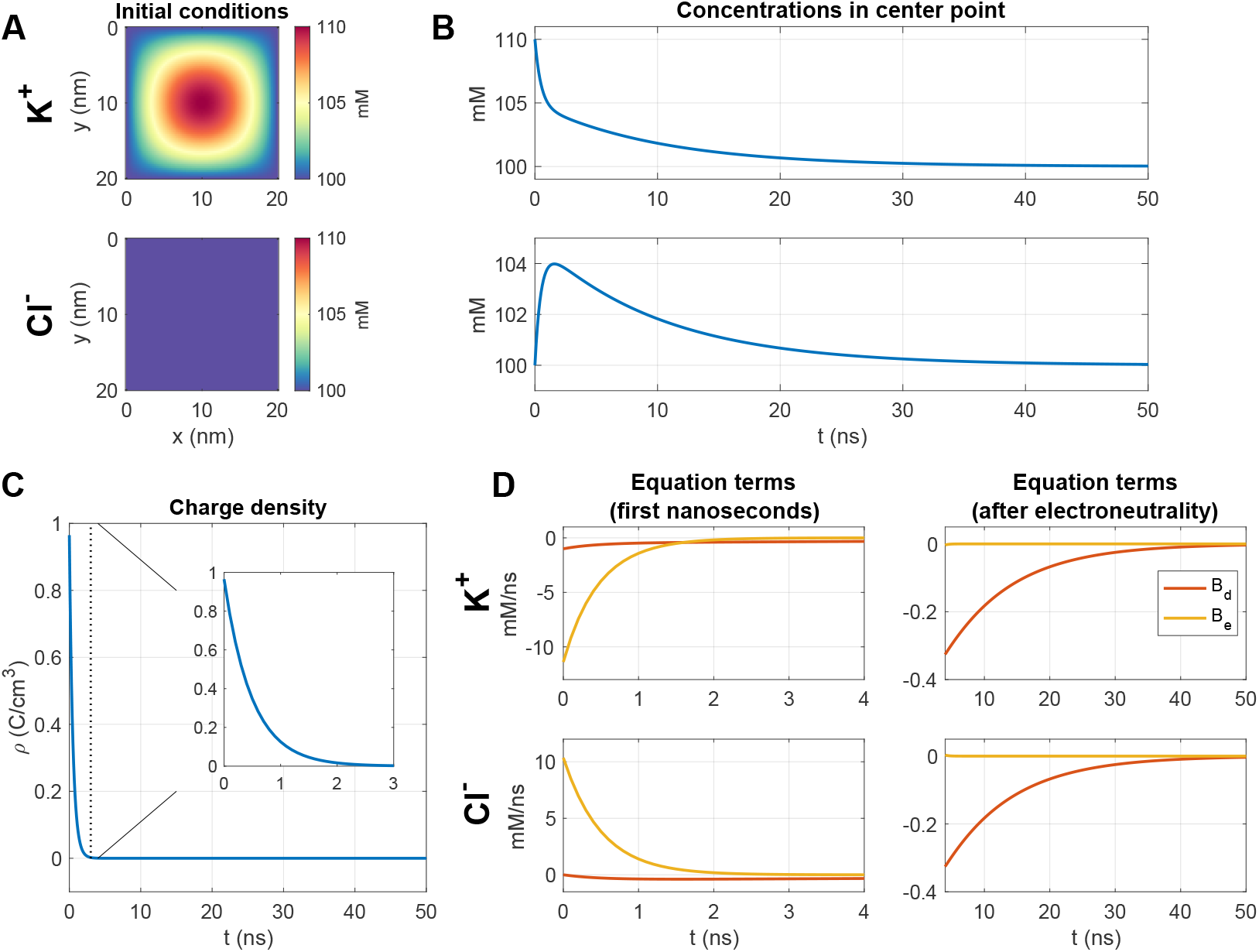
Decay following a perturbation of the K^+^ concentration in a PNP simulation. We consider a simple 2D example with two ions, K^+^ and Cl^*−*^. A: Initial conditions for the two ions. B: Concentrations in the center point as functions of time. C: Charge density, *ρ*, in the center point as a function of time. D: The equation terms, *B*_*d*_ = ∇ · *D*_*k*_∇*c*_*k*_ and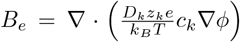, of (2) in the center point as functions of time during the first 4 ns of simulation (left) and after 4 ns (right). We use *D*_K+_ =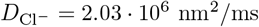, Δ*t* = 0.1 ns and a uniform mesh with Δ*x* = Δ*y* = 0.25 nm.

Figure 5B shows the concentration solutions in the point in the center of the domain for the two ionic species as functions of time. We observe that for the first few nanoseconds, the dynamics are fast and that the dynamics are slower in the later part of the simulation. Figure 5C similarly shows the charge density, *ρ* (defined in (3)), in the center point as a function of time. Here, we observe that the solutions approach electroneutrality (*ρ* = 0) fast during the first 3 ns of the simulation.

In Figure 5D we show the magnitude of the two terms *B*_*d*_ = ∇· *D*_*k*_∇*c*_*k*_ and 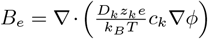 in (2) in the center point for the first few nanoseconds (left panel) and later in the simulation (right panel). We observe that during the first few nanoseconds (before electroneutrality is achieved) the electrical term, *B*_*e*_, is much more prominent than the diffusional term, *B*_*d*_. This electrical term drives the system fast towards electroneutrality during the first few nanoseconds of the simulation. The concentration of K^+^ decreases and the concentration of Cl^*−*^ increases. Next, after electroneutrality is achieved, the electrical term, *B*_*e*_, is negligible, and the diffusional term, *B*_*d*_, is most prominent. This terms slowly brings the concentrations towards the constant value of 100 mM.

### 3.2 Formation of a Debye layer near a membrane in a simple PNP model example

A second simple 2D example illustrating properties of the PNP model is displayed in Figure 6. In this case, we introduce a membrane between two domains, Ω_*L*_ and Ω_*R*_ (see Figure 6C). In the membrane, the diffusion coefficient is set to zero. We again consider the PNP system (1)–(3) for two ions, K^+^ and Cl^*−*^. We use homogeneous Neumann boundary conditions on all boundaries for both the electrical potential and the concentrations, except that we use the Dirichlet boundary condition *ϕ* = 0 mV for the potential on the left boundary. The initial concentration is set to 100 mM for Cl^*−*^ in both Ω_*L*_ and Ω_*R*_, but for K^+^, it is set to is 100.1 mM in Ω_*L*_ and 99.9 mM in Ω_*R*_. This entails that *ρ* is initially slightly positive in Ω_*L*_ and slightly negative in Ω_*R*_. In Figure 6A, we show the solutions near the membrane as functions of *x* for five different points in time. We observe that a layer of adjusted concentrations and non-zero *ρ* gradually forms on each side of the membrane. This layer near the membrane is often referred to as the Debye layer. In Figure 6B, we plot *ρ* as a function of time in one point near the membrane and one point 20 nm right of the membrane. These points are marked as *x*_1_ and *x*_2_ in Figure 6C. We observe that away from the membrane, the solutions approach electroneutrality (i.e., *ρ* approaches zero). Near the membrane, on the other hand, the magnitude of *ρ* increases with time for the first few nanoseconds and then remains constant with time after the Debye layer has been formed.

**Figure 6.**
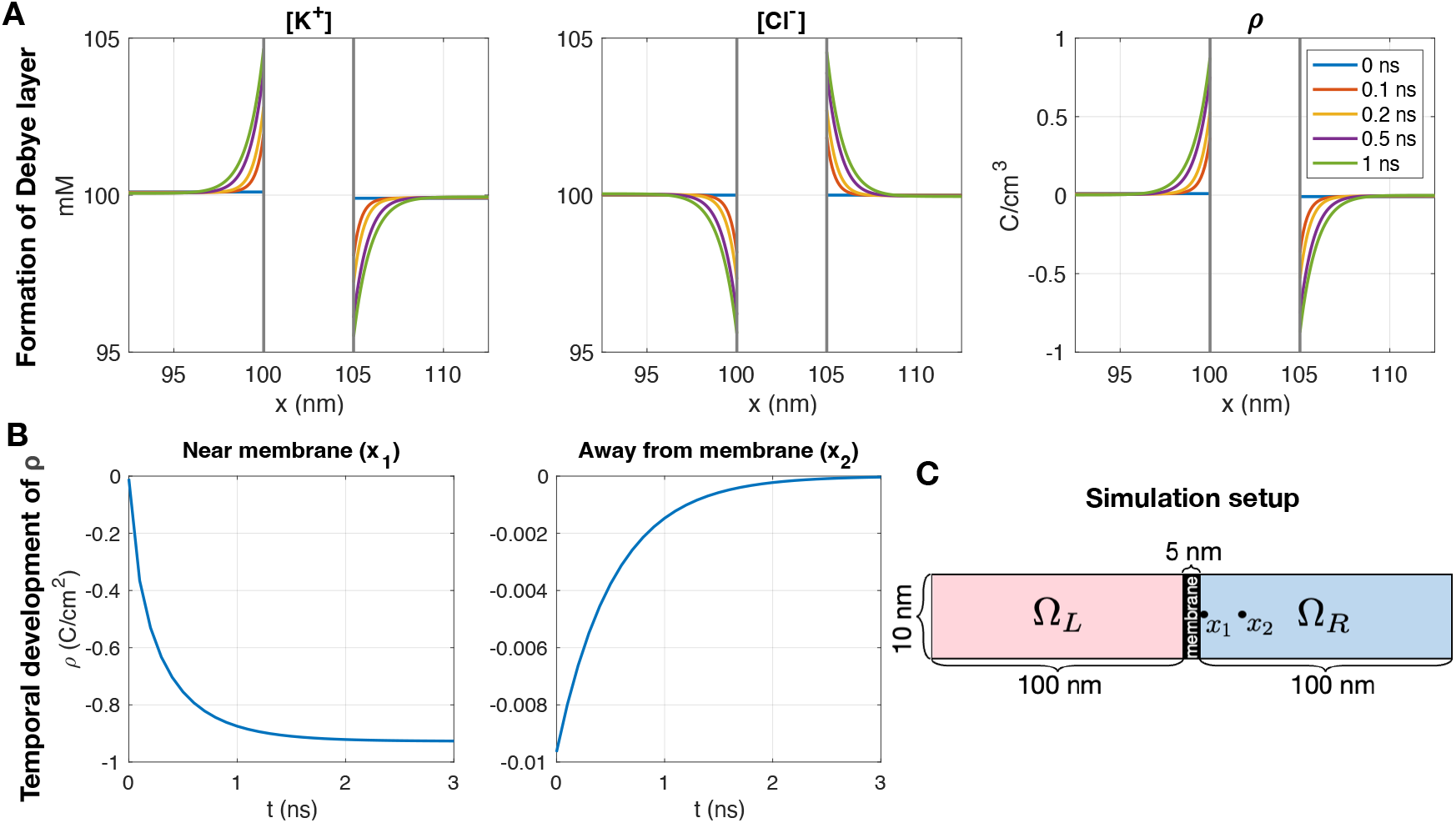
Formation of a Debye layer near a membrane in a PNP simulation. We consider a simple 2D example with two ions. Initially, the concentrations are constant in Ω_*L*_ and Ω_*R*_. The initial concentration of Cl^*−*^ is 100 mM in both Ω_*L*_ and Ω_*R*_, and the initial concentration of K^+^ is 100.1 mM in Ω_*L*_ and 99.9 mM in Ω_*R*_. A: Ionic concentrations and charge density near the membrane as functions of *x* for five different points in time. B: Charge density, *ρ*, 0.1 nm right of the membrane (*x*_1_) and 20 nm to the right of the membrane (*x*_2_) as functions of time. C: Illustration of the simulation setup. We use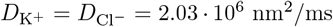, Δ*t* = 0.1 ns and a uniform mesh with Δ*x* = Δ*y* = 0.1 nm.

### 3.3 PNP simulation of an open potassium channel. From electroneutrality to the cell’s resting state

After considering the simple examples in Figures 5 and 6, we will now consider simulations of the full three-dimensional (3D) setup described in the Methods section (see Figure 1). In these simulations, we include intracellular Ca^2+^ binding buffer proteins, i.e., we solve the system (5)–(8).

We first consider the case when only a K^+^ channel is open in the cell membrane. This simulation is used to obtain the resting state of the cell, used as initial conditions for the following simulations. The results of the simulation are displayed in Figure 7. In this visualization, we consider the solutions in a part of the domain that is close to the K^+^ channel. Note that the extracellular part of the domain is depicted on the left-hand side and the intracellular part of the domain is depicted on the right-hand side. Details on the visualization setup are provided in Section 2.6.

**Figure 7.**
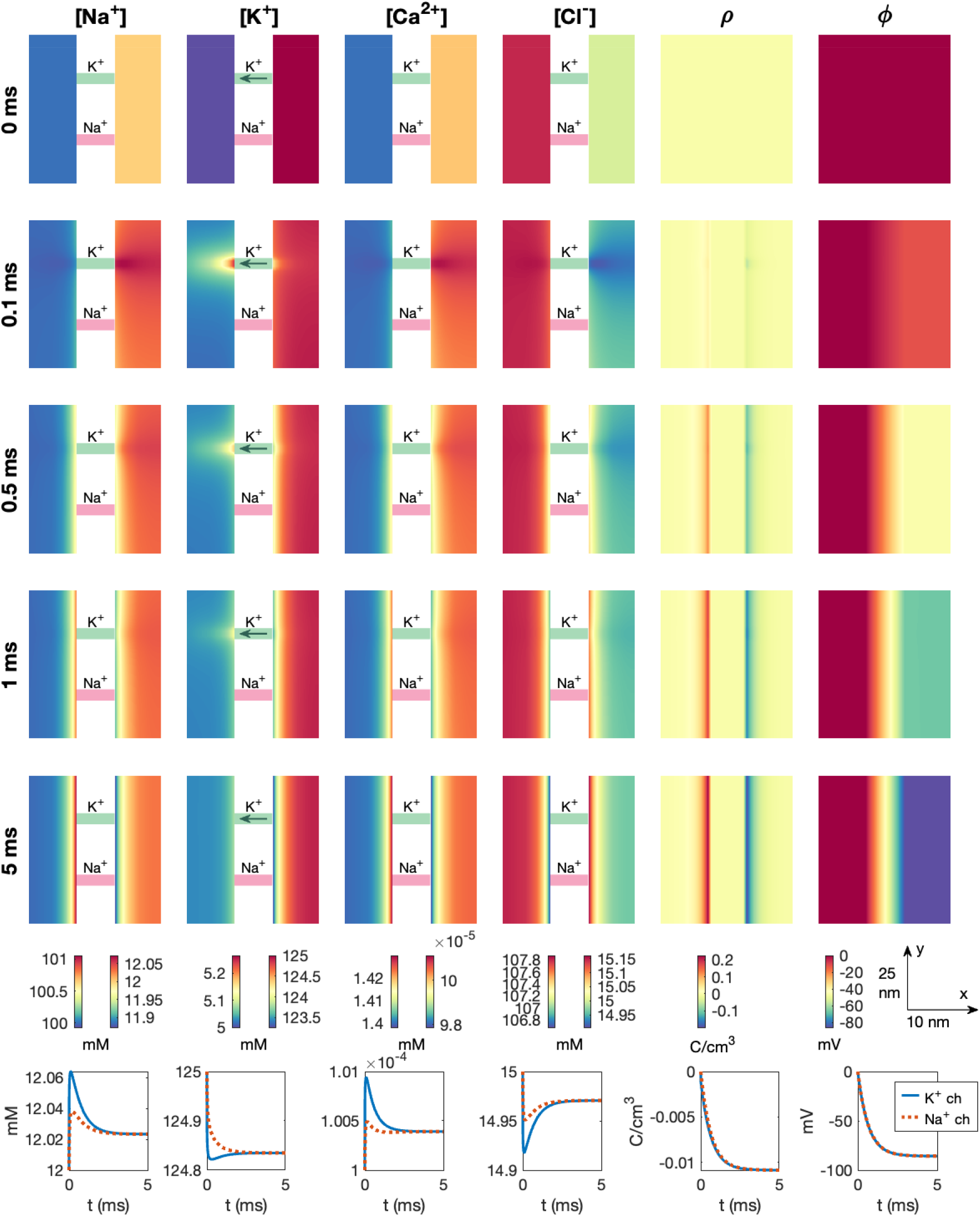
Dynamics following the opening a K^+^ channel in a PNP model simulation. The figure setup is described in Section 2.6. We use Δ*t* = 10 *µ*s and an adaptive mesh like illustrated in Figure 3.

In Figure 7, we observe that, initially, all ionic concentrations are constant in the intracellular and extracellular parts of the domain (but different in these two parts of the domain). Moreover, the system fulfills electroneutrality (*ρ* = 0) and the electrical potential is zero everywhere. As the K^+^ channel is opened, we observe that K^+^ ions move from the intracellular to the extracellular part of the domain. This results in an elevated extracellular K^+^ concentration and a reduced intracellular K^+^ concentration, especially in the vicinity of the K^+^ channel (see, e.g., *t* = 0.1 ms). As was observed in Figure 5, the resulting deviation from electroneutrality near the channel is counteracted by local changes in the other ionic concentrations. For example, we observe that the concentration of the positive ions (Na^+^ and Ca^2+^) increases slightly on the intracellular side of the K^+^ channel, whereas the concentration of the negative Cl^*−*^ ions oppositely decreases. These effects act to counteract the reduced K^+^ concentration on the intracellular side of the channel.

Nevertheless, since no other ions than K^+^ is able to cross the membrane, the movement of K^+^ from the intracellular to the extracellular side of the membrane ultimately leads to a surplus of positive charges in the extracellular space and a surplus of negative charges in the intracellular space. As was also observed in Figure 6, this deviation from electroneutrality on opposite sides of the membrane results in a layer of positive charge density, *ρ*, on the extracellular side of the membrane and a layer of negative charge density on the intracellular side of the membrane (i.e., a Debye layer). For the positive ions (Na^+^, K^+^, and Ca^2+^), this entails a layer of increased concentration on the extracellular side of the membrane and a layer of decreased concentration of the intracellular side. Conversely, a layer of decreased Cl^*−*^ concentration forms on the extracellular side of the membrane and a layer of increased Cl^*−*^ concentration forms on the intracellular side (see, e.g., *t* = 5 ms). Moreover, a potential difference is generated over the membrane. After a steady state solution is formed, the transmembrane potential, *v* = *ϕ*_*i*_ − *ϕ*_*e*_, is about −80 mV, determined by the Nernst equilibrium potential of K _^+^_ (see (12)).

In Figure 8, the solutions near the membrane at the end of the simulation is displayed, showing the details of the obtained Debye layer.

**Figure 8.**
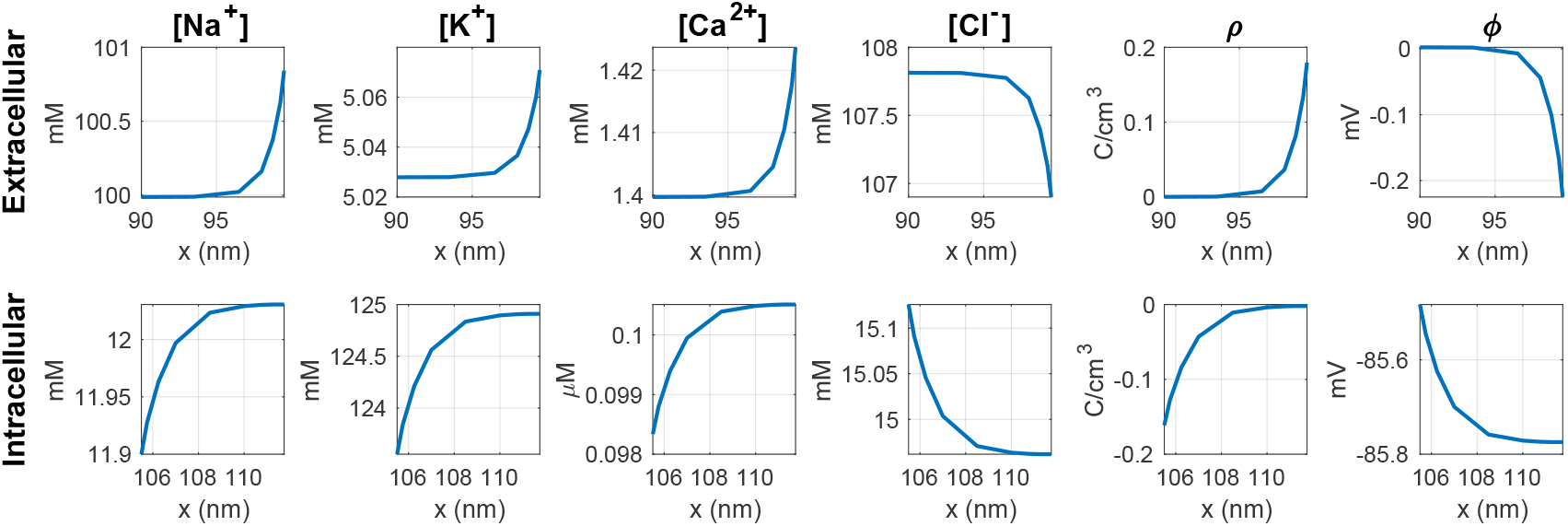
Steady state solutions close to the membrane for the cell at rest. Only a K^+^ channel is open. The plots show the solutions along a line in the *x*-direction, taken from the simulation shown in Figure 7.

**Figure 9.**
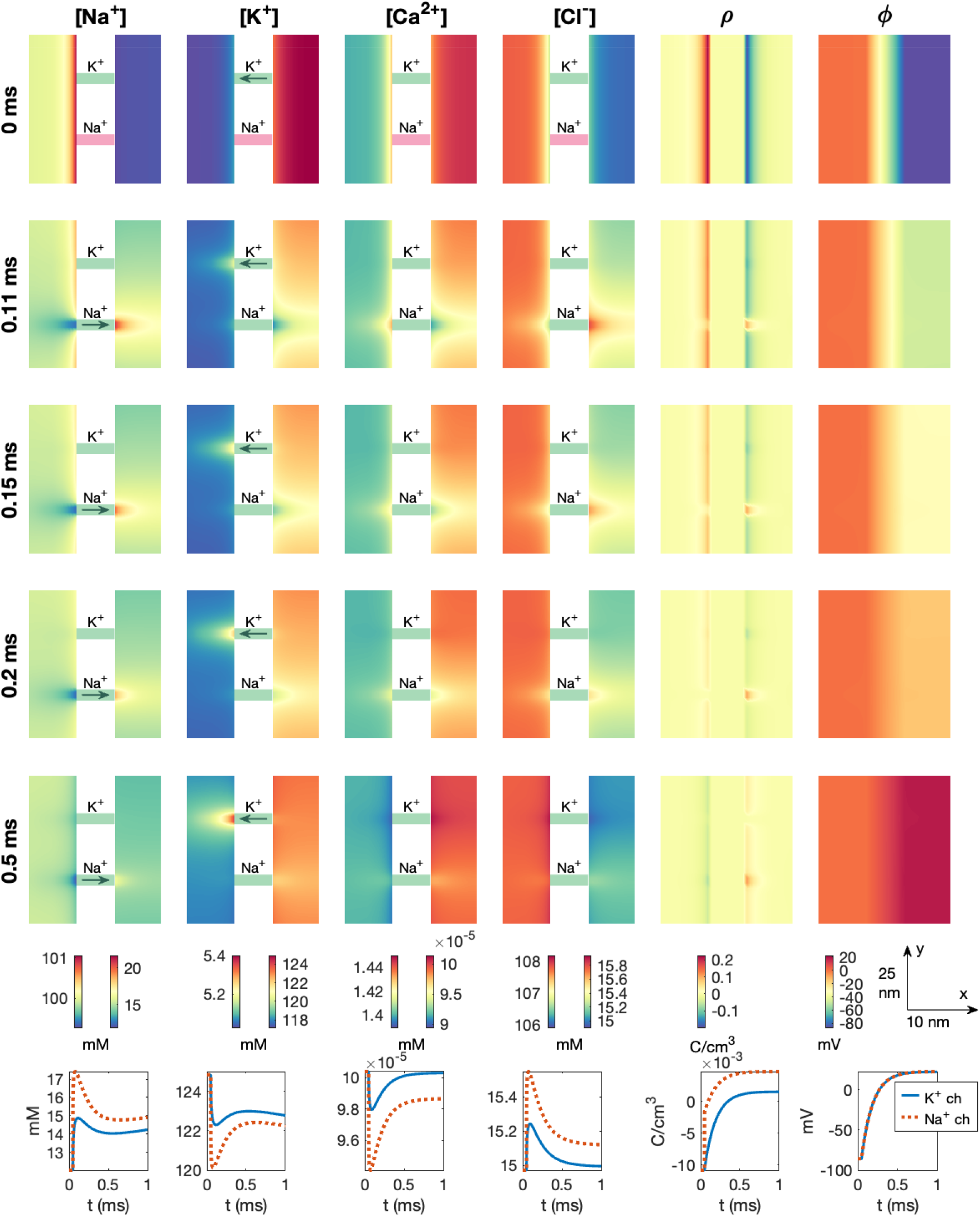
Dynamics following the opening of a Na^+^ channel in a PNP model simulation. The figure setup is described in Section 2.6. We use Δ*t* = 1 *µ*s and an adaptive mesh like illustrated in Figure 3.

### 3.4 PNP simulation of an open sodium channel

Next, we consider the case of opening a Na^+^ channel to generate an action potential upstroke. We start from the initial conditions representing the cell’s resting state, taken from the end of the simulation displayed in Figure 7. We keep the K^+^ channel open and open the Na^+^ channel at *t* = 0.05 ms. Figure 9 shows the results of this simulation. We observe that after the Na^+^ channel is opened, Na^+^ moves from the extracellular to the intracellular space, resulting in a locally increased Na^+^ concentration on the intracellular side of the Na^+^ channel and a locally decreased Na^+^ concentration on the extracellular side (see, e.g., *t* = 0.11 ms). To counteract the local deviation from electroneutrality, the concentration of the other ionic species also change locally in the vicinity of the Na^+^ channel. For example, the K^+^ and Ca^2+^ concentrations decrease on the intracellular side of the Na^+^ channel, and the Cl^*−*^ concentration increases.

Moreover, the Na^+^ ions moving from the extracellular to the intracellular domain, gradually reduce the surplus of positive charges in the extracellular space and the surplus of negative charges in the intracellular space, reducing the magnitude of *ρ* near the membrane. In fact, eventually, there is a surplus of *negative* charges on the extracellular side of the membrane and a surplus of *positive* charges on the intracellular side (see *t* = 0.5 ms). The intracellular potential (and thus also the transmembrane potential) increases from about −80 mV to about 20 mV during the course of the simulation. This upstroke lasts for about 0.5 ms (see rightmost lower panel of Figure 9).

We also note that as the transmembrane potential is increased from the Nernst equilibrium potential for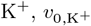, (see (14)) we also get current through the K^+^ channel, which leads to local concentration changes close to the K^+^ channel.

### 3.5 PNP simulation of an open calcium channel

After observing the dynamics following the opening of a K^+^ channel leading to the resting state of the cell (Figure 7) and the opening of a Na^+^ channel leading to the action potential upstroke (Figure 9), we will now consider the dynamics taking place in the dyad. We first consider the case of opening a Ca^2+^ channel. More specifically, we start the simulation at the resting state with only a K^+^ channel open. This K^+^ channel remains open during the entire simulation. At *t* = 0.05 ms, the Na^+^ channel is opened (like in Figure 9) starting the initial phase of the action potential upstroke. Subsequently, at *t* = 0.1 ms, we open the Ca^2+^ channel in the dyad. The results of this simulation is shown in Figure 10, and in this case the visualization focuses on the area close to the Ca^2+^ channel and NCX (i.e, the dyad).

**Figure 10.**
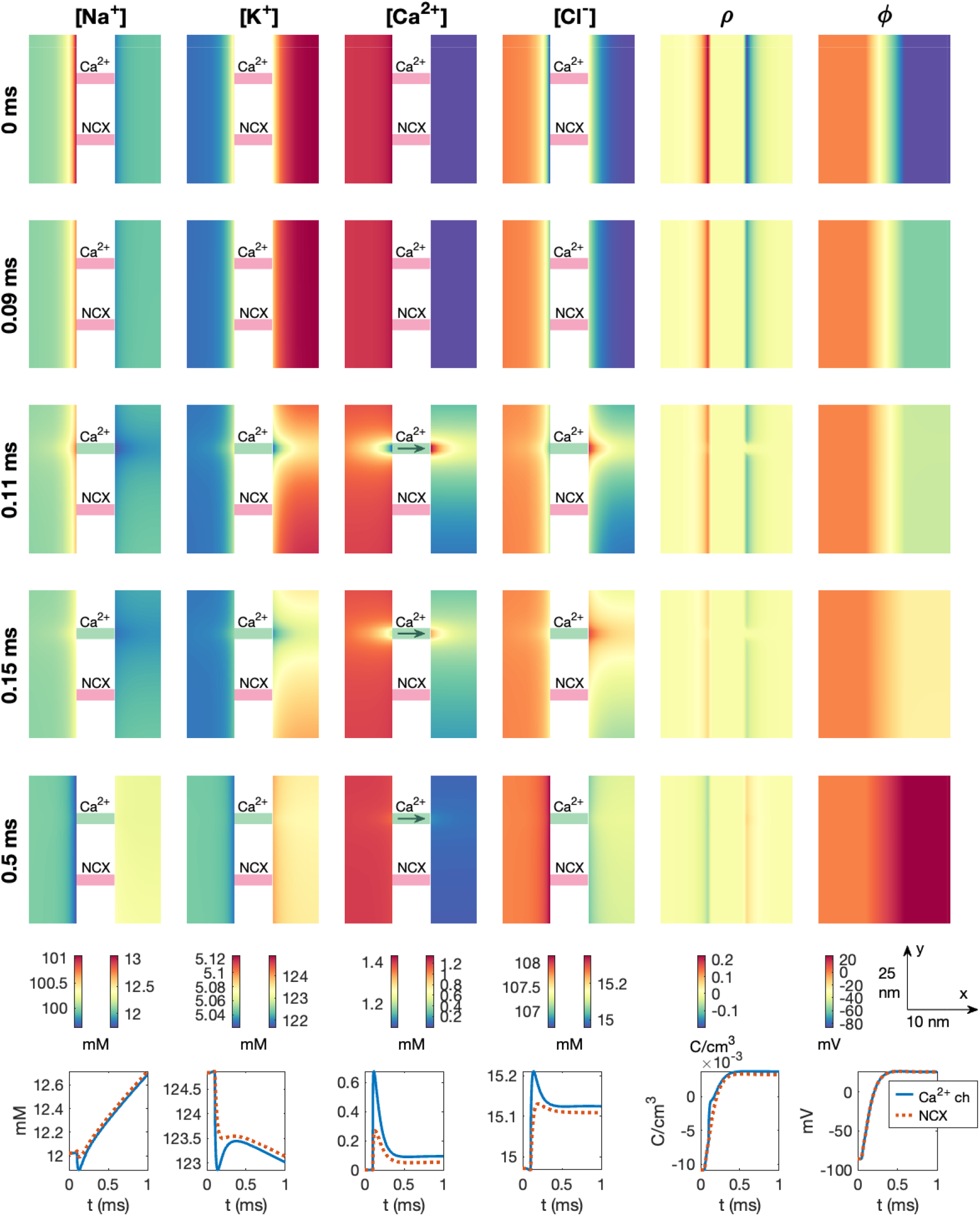
Dyad dynamics following the opening of a Ca^2+^ channel in a PNP model simulation. The figure setup is described in Section 2.6. We use Δ*t* = 1 *µ*s and an adaptive mesh like illustrated in Figure 3.

We observe that a Debye layer is present in the dyad. Moreover, at *t* = 0.09 ms, following the Na^+^ channel opening at *t* = 0.05 ms, the magnitude of *ρ* near the membrane is reduced and the intracellular potential (and thus the transmembrane potential) is increased somewhat compared to the resting state (*t* = 0 ms). After the Ca^2+^ channel is opened at *t* = 0.1 ms, the Ca^2+^ concentration in the dyad increases, especially in the vicinity of the Ca^2+^ channel. However, there is a clearly visible increase in the Ca^2+^ concentration across the width of the dyad as well. In the lower panel, we observe that halfway across the dyad outside of the Ca^2+^ channel, the Ca^2+^ concentration increases from 0.0001 mM to about 0.7 mM. Further away in the *y*-direction, halfway across the dyad outside of the closed NCX, the concentration increases to about 0.3 mM. This increased Ca^2+^ concentration could trigger RyR opening on the membrane of the SR. As observed in Figures 7 and 9, we also observe local changes in the remaining ionic species close to the Ca^2+^ channel, counteracting the deviation from electroneutrality resulting from the Ca^2+^ influx.

### 3.6 PNP simulation with a dyad including a sodium calcium exchanger and an open calcium channel

In Figure 11, we show the result of a simulation similar to the one displayed in Figure 10. The difference from Figure 10 is that the NCX is open, and it remains open during the entire simulation. Considering the arrows showing the direction of flow through the NCX, we observe that at rest, the NCX transports Ca^2+^ out of the cell and Na^+^ into the cell, but following the opening of the Na^+^ channel, the flux changes direction (see *t* = 0.09 ms). After the Ca^2+^ channel is opened, however, the direction changes back to transporting Ca^2+^ out of the cell and Na^+^ into the cell. Nonetheless, comparing the solutions of Figure 10 and Figure 11, the presence of an open NCX does not appear to significantly change the dyad solutions. For instance, the Ca^2+^ concentration in the points halfway across the dyad plotted in the lower figure panels appears to be very similar regardless of the presence of an open NCX.

**Figure 11.**
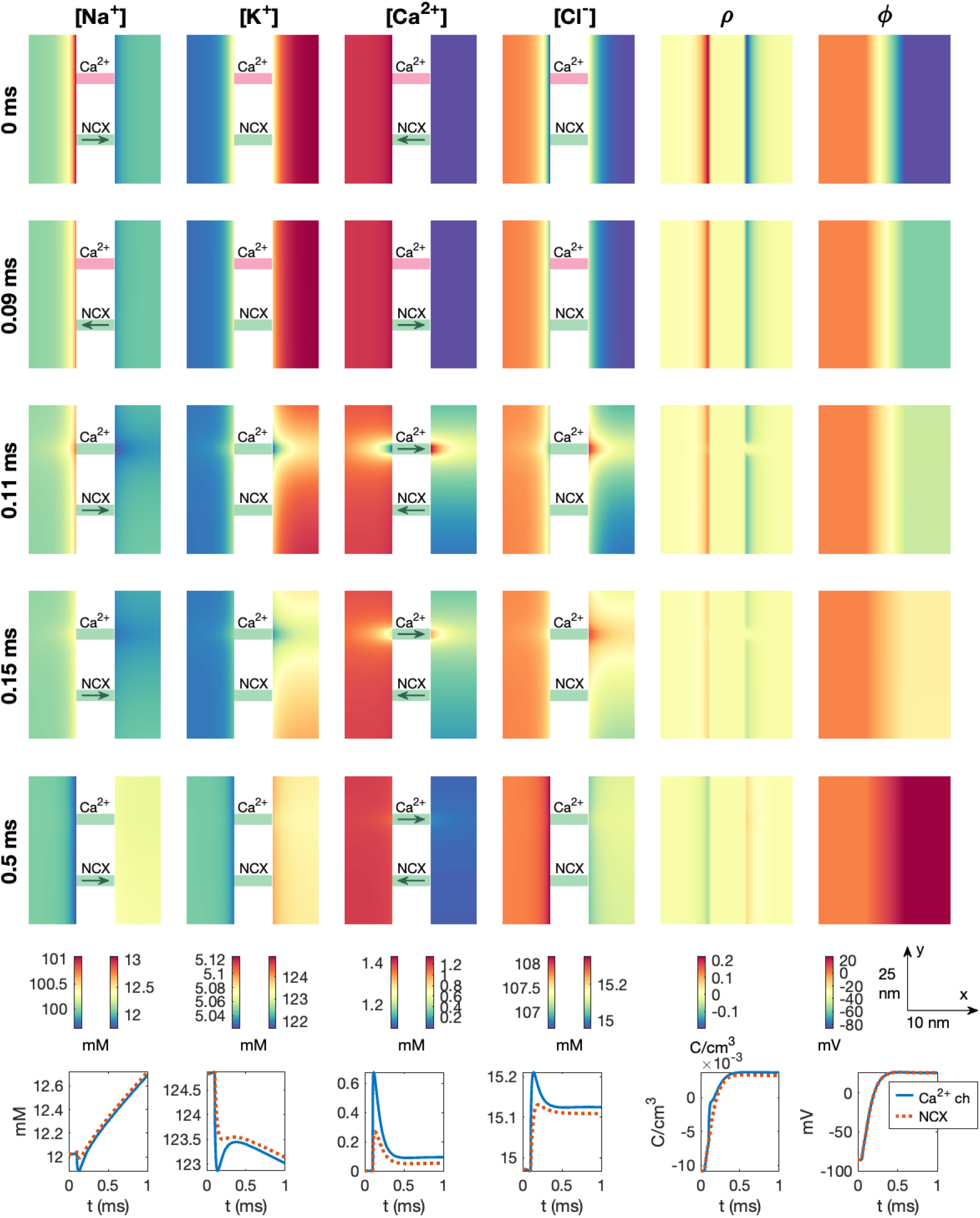
Dyad dynamics following the opening of a Ca^2+^ channel in a PNP model simulation including an open NCX. The figure setup is described in Section 2.6. We use Δ*t* = 1 *µ*s and an adaptive mesh like illustrated in Figure 3.

### 3.7 PNP simulation of a dyad with a sodium calcium exchanger and no open calcium channel

To demonstrate a potential effect of an open NCX present in the dyad, we consider the case when the Ca^2+^ channel is closed. The K^+^ channel is open during the entire simulation, and the Na^+^ channel is opened at *t* = 0.05 ms. The results of this simulation is displayed in Figure 12. We observe that in this case, the NCX flux remains in the direction that transports Ca^2+^ ions into the cell after the Na^+^ channel opening. At *t* = 1 ms, this results in a significant influx of Ca^2+^ into the dyad through the NCX. Halfway across the dyad, the concentration reaches about 2·10^*−*4^ mM = 0.2 *µ*M. This increase in Ca^2+^ concentration could potentially affect RyRs located in the apposing SR membrane.

**Figure 12.**
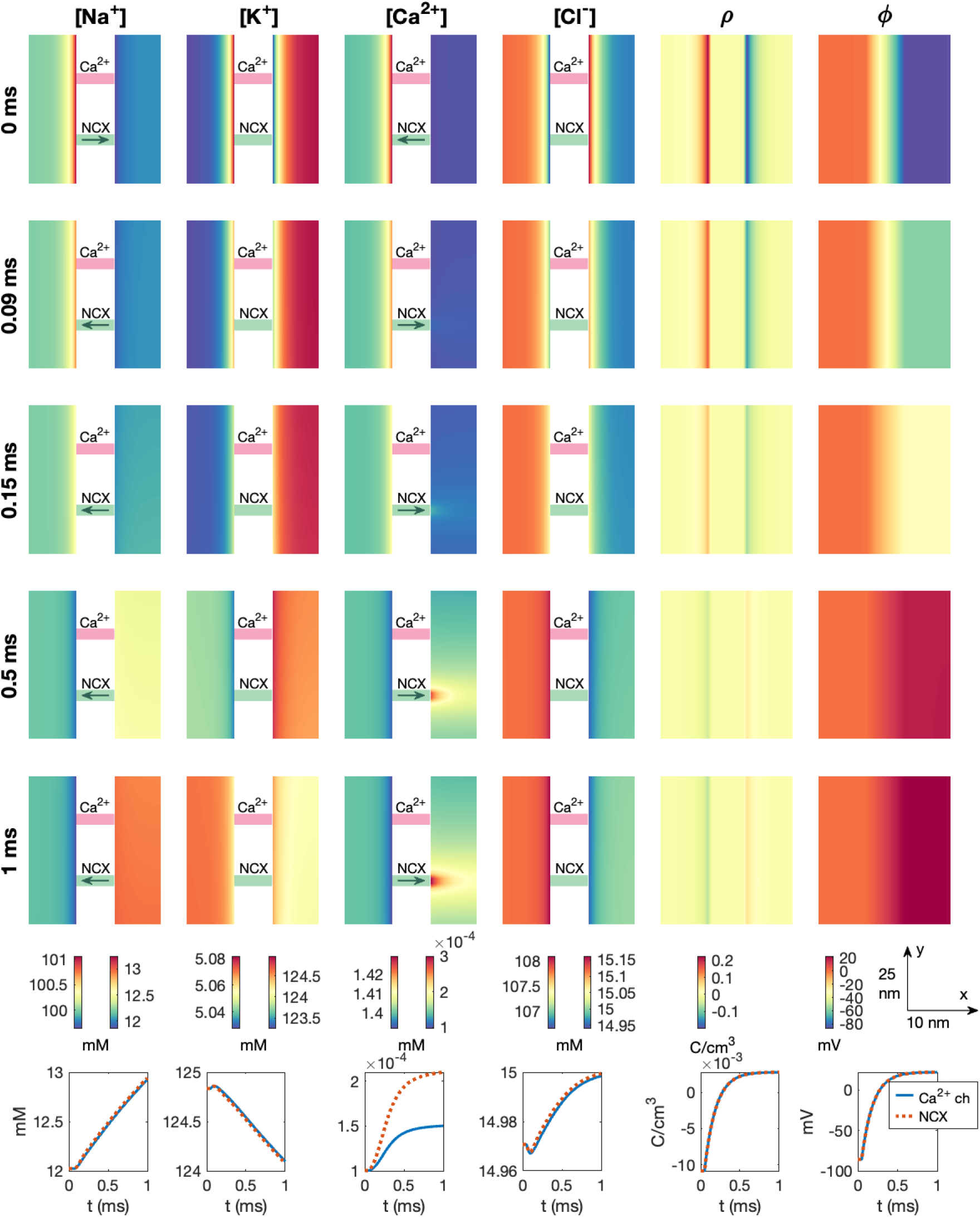
Dyad dynamics following the opening of an Na^+^ channel in a PNP model simulation including an NCX. The Ca^2+^ channel is not opened. The figure setup is described in Section 2.6. We use Δ*t* = 1 *µ*s and an adaptive mesh like illustrated in Figure 3.

### 3.8 PNP simulation with more prominent effects of the sodium calcium exchanger

To increase the effects of the NCX flux, we also consider a case in which the dyad width, *L*_*i*_, is decreased to 5 nm, all intracellular diffusion coefficients are reduced to half of the default values reported in Table 1, and the Na^+^ channel is moved to be in close proximity of the NCX. These changes are introduced in an attempt to increase the effect of the NCX flux in the dyad. We still keep the Ca^2+^ channel closed. The results of this simulation is shown in Figure 13. We observe that the effect of the NCX flux is larger in this case, and the Ca^2+^ concentration reaches a value of about 0.45 *µ*M halfway across the dyad at *t* = 1 ms.

**Figure 13.**
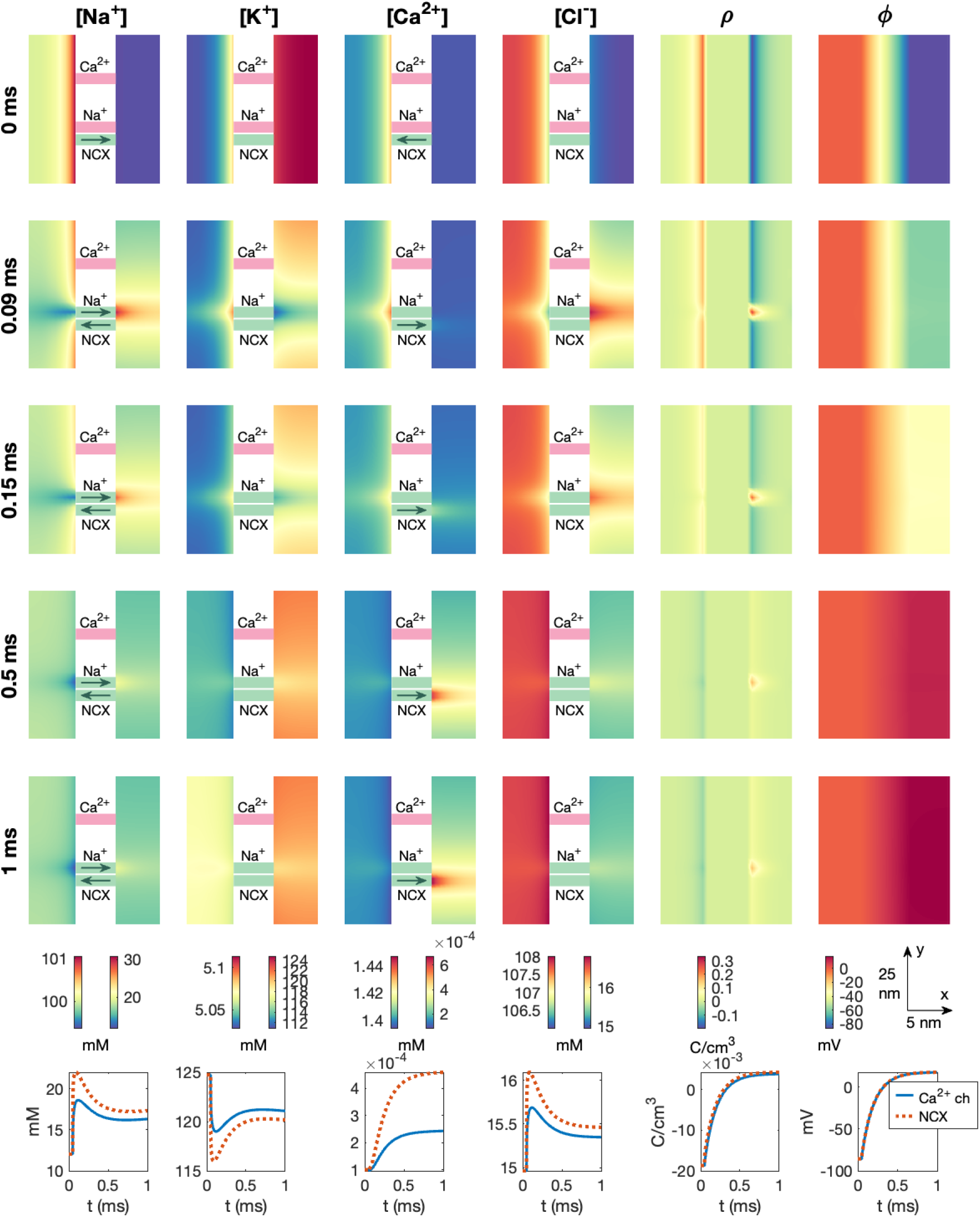
Dyad dynamics following the opening of a Na^+^ channel near an NCX in a PNP model simulation. The Ca^2+^ channel is not opened. The figure setup is described in Section 2.6. We use Δ*t* = 1 *µ*s and an adaptive mesh like illustrated in Figure 3.

### 3.9 Investigating the effect of the dyad width and diffusion coefficient

In Figure 11, we presented the dyad dynamics in the default case of an open Ca^2+^ channel and an NCX present in the dyad. We observed that following the Ca^2+^ channel opening at *t* = 0.1 ms, the Ca^2+^ concentration in the dyad increased. If RyRs were present at the right side of the dyad, this increased Ca^2+^ concentration could have triggered the opening of RyRs on the membrane of the SR.

In Figure 14, we investigate how long it takes from the Ca^2+^ channel is opened in the cell membrane until the RyRs would be triggered. We refer to this duration as the RyR activation time. The RyR activation times reported in Figure 14 are defined as the duration from the Ca^2+^ channel is opened at *t* = 0.1 ms until the point in time when the average Ca^2+^ concentration in an area corresponding to an RyR (i.e., an area spanning 30 nm × 30 nm directly across the dyad from the Ca^2+^ channel) reaches a value above 0.5 *µ*M, representing the threshold for RyR activation [34].

**Figure 14.**
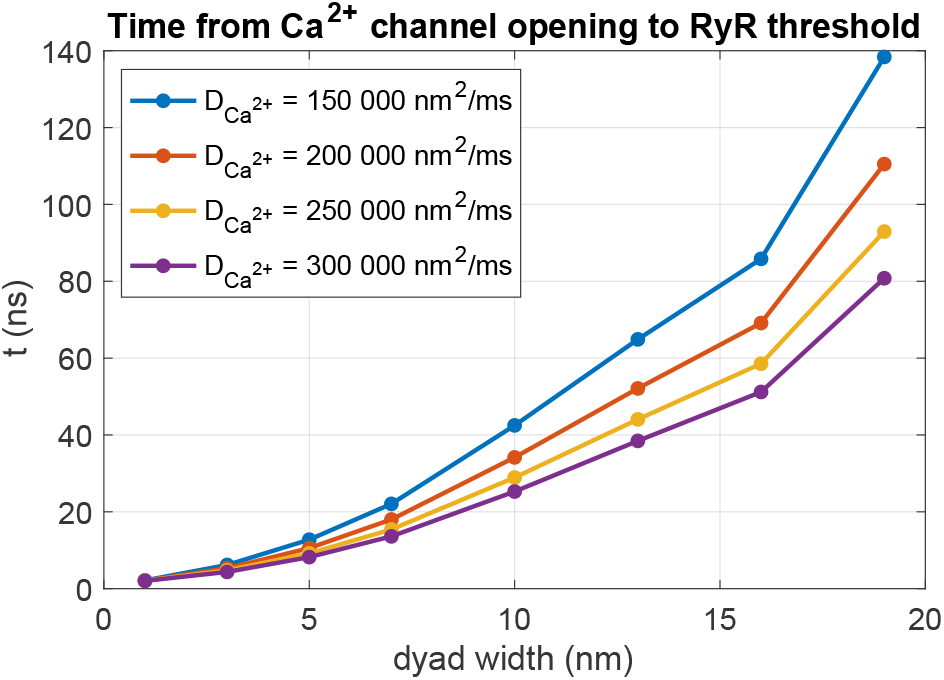
RyR activation time for different values of the dyad width (*L*_*i*_) and the intracellular Ca^2+^ diffusion coefficient (*D*_Ca2+_). The RyR activation time is defined as time from the membrane Ca^2+^ channel is opened until the Ca^2+^ concentration outside of an apposing RyR channel reaches 0.5 *µ*M. We perform PNP model simulations similar to the one displayed in Figure 11. We use Δ*t* = 1 ns and an adaptive mesh like illustrated in Figure 3.

We investigate how the RyR activation time depends on the dyad width (*L*_*i*_) and the intracellular Ca^2+^ diffusion coefficient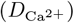. We consider dyad widths between 1 nm and 19 nm. Note that for dyad widths below 5 nm, we refine the resolution near the membrane from the default Δ*x* = 0.5 nm to Δ*x* = 0.25 nm. Additionally, we consider values of the intracellular Ca^2+^ diffusion coefficient between 150 000 nm^2^/ms and 300 000 nm^2^/ms, based on different values applied for Ca^2+^ diffusion in the dyad found in literature [37, 38, 28, 39, 40, 41]. The intracellular diffusion coefficients for the remaining ionic species provided in Table 1 are scaled by the same factor as for Ca^2+^.

Figure 14 shows that the RyR activation time increases in a non-linear manner as the dyad width is increased. Moreover, the RyR activation time increases as the diffusion coefficient is decreased.

### 3.10 Comparison to the pure diffusion model

We will now compare the above reported solutions of the PNP model to the solution of a pure diffusion or reaction-diffusion model.

#### 3.10.1 Decay following perturbations

We start the comparison between the PNP and diffusion models by considering the simple example of an initial perturbation displayed in Figure 5. In Figure 15, we compare the solution of the full PNP system (1)–(3) to the solution of the pure diffusion equation

**Figure 15.**
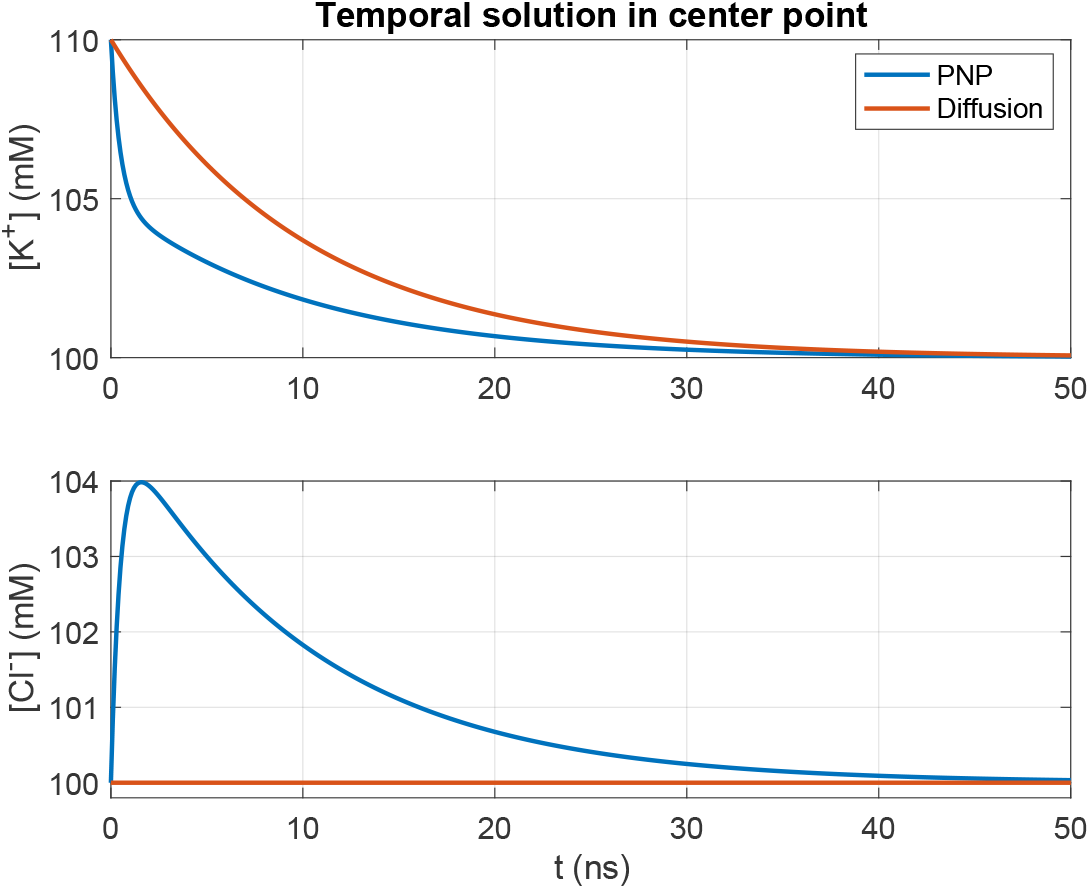
Decay following a perturbation of one of the concentrations in simulations of the PNP model and the pure diffusion model. We consider the simple 2D example with two ions displayed in Figure 5. The initial conditions of the two ions are shown in Figure 5A, and we plot the concentrations in the point in the center of the domain, like in Figure 5B. We have used Δ*t* = 0.1 ns and a uniform mesh with Δ*x* = Δ*y* = 0.25 nm.

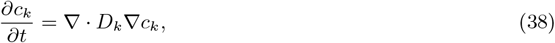

for *k* = {K^+^, Cl^*−*^}. For the pure diffusion model, there is no equation terms driving the system towards electroneutrality. Therefore, the Cl^*−*^ concentration is unaffected by the perturbation in the K^+^ concentration and remains constant throughout the simulation. Furthermore, there are no initial fast dynamics driving the K^+^ concentration towards electroneutality. Thus the decay of the K^+^ concentration is slower for the pure diffusion model compared to the PNP model.

#### 3.10.2 Debye layer near a membrane

We next consider the case of the Debye layer formation near a membrane investigated in Figure 6. Since the concentrations are constant on each side of the membrane and the membrane do not allow for diffusion, the right-hand side of the pure diffusion model (38) would be zero for the setup used as initial conditions for this example. Thus, the concentrations would remain constant, and no Debye layer would be formed in a pure diffusion model simulation. In other words, the PNP model is required to represent the Debye layer.

#### 3.10.3 Dyad dynamics

We now consider the dyad simulation setup (see Figure 1C) and compare the solution of the full PNP system (5)–(8) to the solution of the reaction-diffusion equations

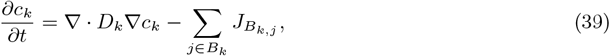

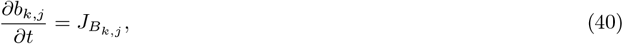

where 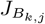 is defined in (4). We consider the same buffers and parameters as in the PNP model simulations (see Table 2).

In Figure 16, we show the solution of a reaction-diffusion model simulation of the same setup and open channels as in the PNP model simulation displayed in Figure 11. Since the reaction-diffusion system (39)–(40) do not give rise to an electrical potential, we have used the potential computed in the PNP model simulation to compute the transmembrane fluxes. Like in Figure 15, we observe that for the reaction-diffusion model a local increase in the Ca^2+^ concentration do not affect any of the other ionic species like it does in the PNP model (compare Figures 16 and 16). In addition, no Debye layer is present for the reaction-diffusion model. Therefore, the concentrations of all ionic species except for Ca^2+^ are constant in the dyad, unlike in the PNP simulation, where local gradients are present for all ions near the membrane. Nevertheless, the Ca^2+^ concentration outside of the Ca^2+^ channel, on the other hand, appears to be very similar in the reaction diffusion-model and the PNP model simulation (compare Figures 16 and 16).

**Figure 16.**
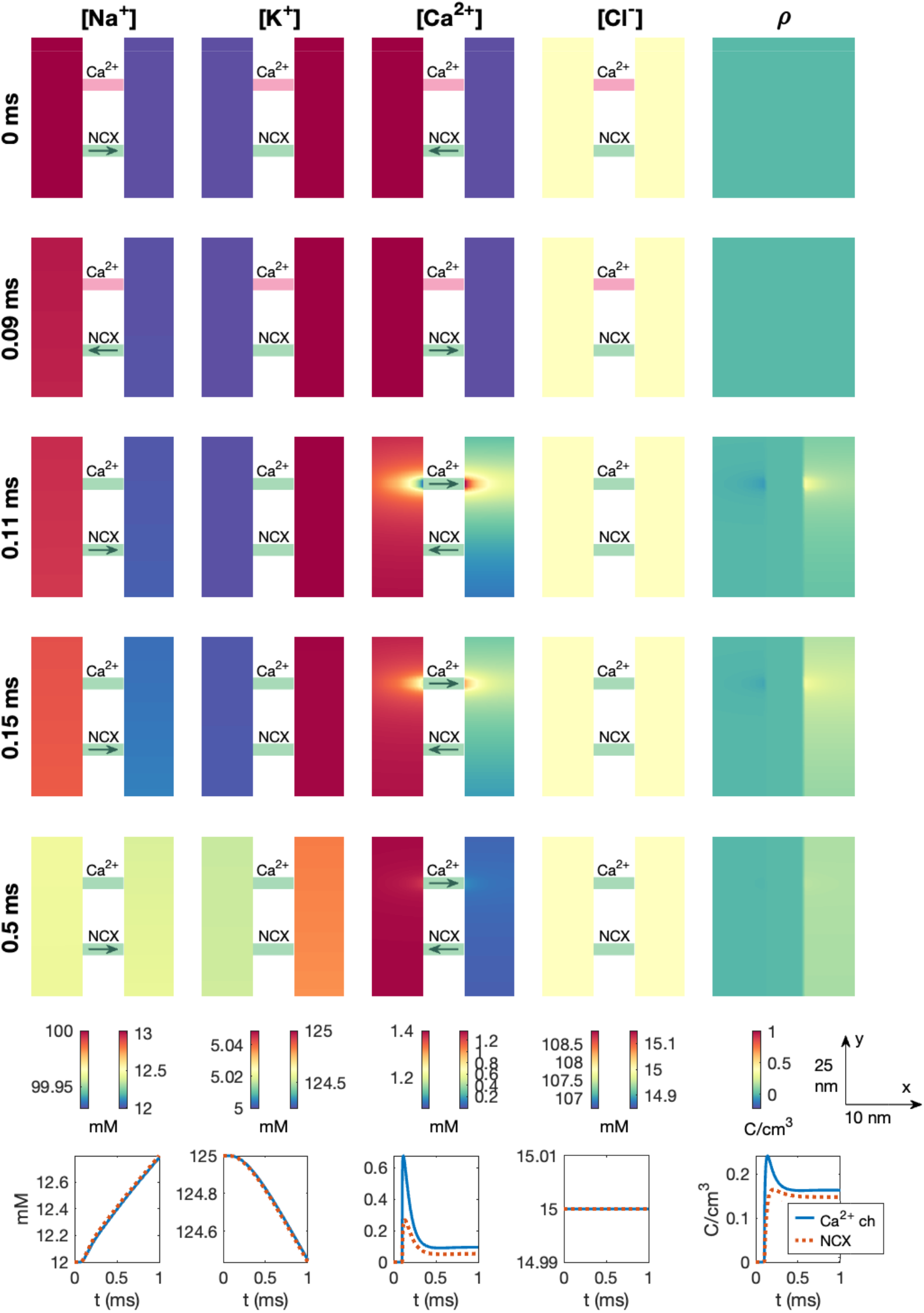
Dyad dynamics following the opening of a Ca^2+^ channel in a pure reaction-diffusion model version of the PNP simulation displayed in Figure 11. The figure setup is described in Section 2.6. We have used Δ*t* = 1 *µ*s and an adaptive mesh like illustrated in Figure 3.

In Figure 17, we investigate whether the differences between the PNP model and the reaction-diffusion model observed in Figures 16 has an effect of the time from Ca^2+^ channel opening until the triggering of an RyR, defined as the RyR activation time. In Figure 17, we observe that the RyR activation time computed using the reaction-diffusion model appears to be virtually identical to that computed using the full PNP model. This also holds as the dyad width or the intracellular diffusion coefficients are adjusted. Figure 17 thus indicates that the reaction-diffusion model might be sufficient for capturing some of the important Ca^2+^ dynamics in the dyad despite the solution differences from the PNP model observed in Figures 15 and 16.

**Figure 17.**
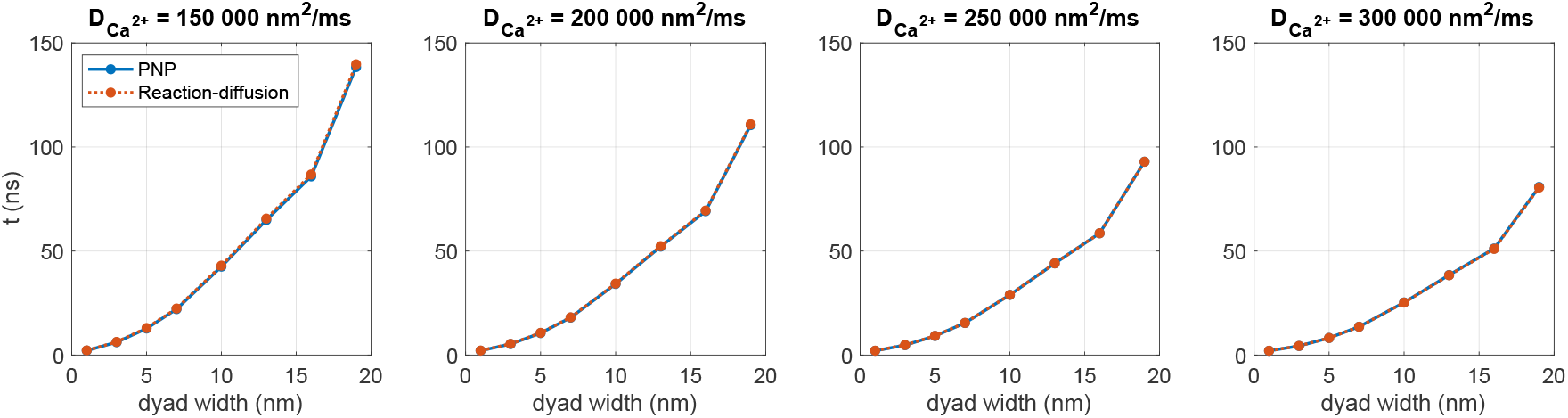
RyR activation time for different values of the dyad width (*L*_*i*_) and the intracellular Ca diffusion coefficient 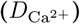 in the PNP and reaction-diffusion models. The RyR activation time is defined as time from the membrane Ca^2+^ channel is opened until the Ca^2+^ concentration outside of an apposing RyR channel reaches 0.5 *µ*M. The PNP results are also displayed in Figure 14. We have used Δ*t* = 1 *µ*s and an adaptive mesh like illustrated in Figure 3. Note that the electrical potential computed in the PNP model is used to compute the transmembrane fluxes in the reaction-diffusion model.

Note that a significant difference between the PNP model and the reaction-diffusion model is that the PNP model gives rise to an electrical potential whereas the reaction-diffusion model does not. If a reaction-diffusion model should be applied to study the Ca^2+^ dynamics in the dyad, the transmembrane potential involved in the transmembrane fluxes would have to be computed in some other manner, e.g., by using a standard ODE model representation of the action potential, as, e.g., done in [34].

### 3.11 Comparison to ODE model representations

We also wish to compare the PNP model of the dyad dynamics to a model that is based on further modeling simplifications. More specifically, we consider an ODE version of the dyad dynamics, similar to what is often applied in action potential models (see, e.g., [42, 43, 29, 35]).

#### 3.11.1 ODE model representation with one dyad compartment

We first consider an ODE model using the most typical representation of the dyad, i.e., by representing the dyad as one computational compartment, Ω_*d*_ (see Figure 18A). In addition, we consider one large surrounding cytosol compartment, Ω_*l*_. The concentrations in these two compartments are represented as constant in space, but varying in time, and we consider Ca^2+^ ions and Ca^2+^ binding buffers. Ca^2+^ ions are allowed to diffuse between the two compartments, represented by the flux *J*_*d,l*_. In addition, Ca^2+^ flows into the dyad compartment through a membrane Ca^2+^ channel (*J*_ch_). The model equations of this ODE model are described in the Supplementary Information.

**Figure 18.**
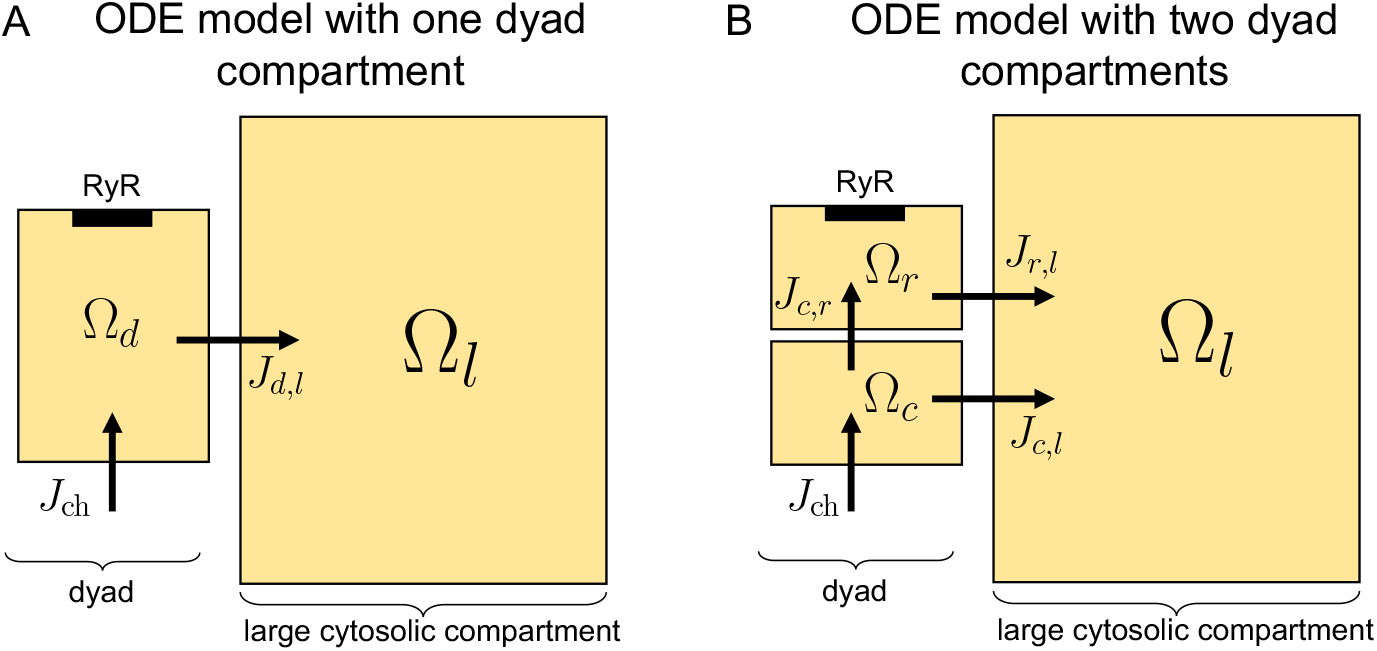
Illustration of two ODE model representations of the dyad dynamics. The ionic concentrations are assumed to be constant (in space) in each considered compartment. A: The ODE model consists of one dyad compartment, Ω_*d*_, and one larger compartment representing the surrounding cytosol, Ω_*l*_. B: The ODE model consists of two dyad compartments (one near the Ca^2+^ channel, Ω_*c*_, and one near the RyR, Ω_*r*_), in addition to the larger compartment representing the surrounding cytosol, Ω_*l*_.

In the leftmost panel of Figure 19, we have performed simulations using this ODE model to identify the RyR activation time, like for the PNP model in Figure 14. Since the dyadic Ca^2+^ concentration, *c*_*d*_, is assumed to be constant in space, this amounts to finding the time it takes from Ca^2+^ channel opening until *c*_*d*_ ≥ 0.5 *µ*M. The parameters of the ODE model are adjusted such that this time is the same as in the PNP model for 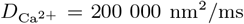 and L*i* = 7 nm. We observe that as the dyad width is increased, the RyR activation time is increased, like also observed for the PNP model (see rightmost panel). However, the ODE model underestimates the increase in time as the dyad width in increased. In addition, altering the diffusion coefficient appears to have very little effect on the RyR activation time in this ODE model.

**Figure 19.**
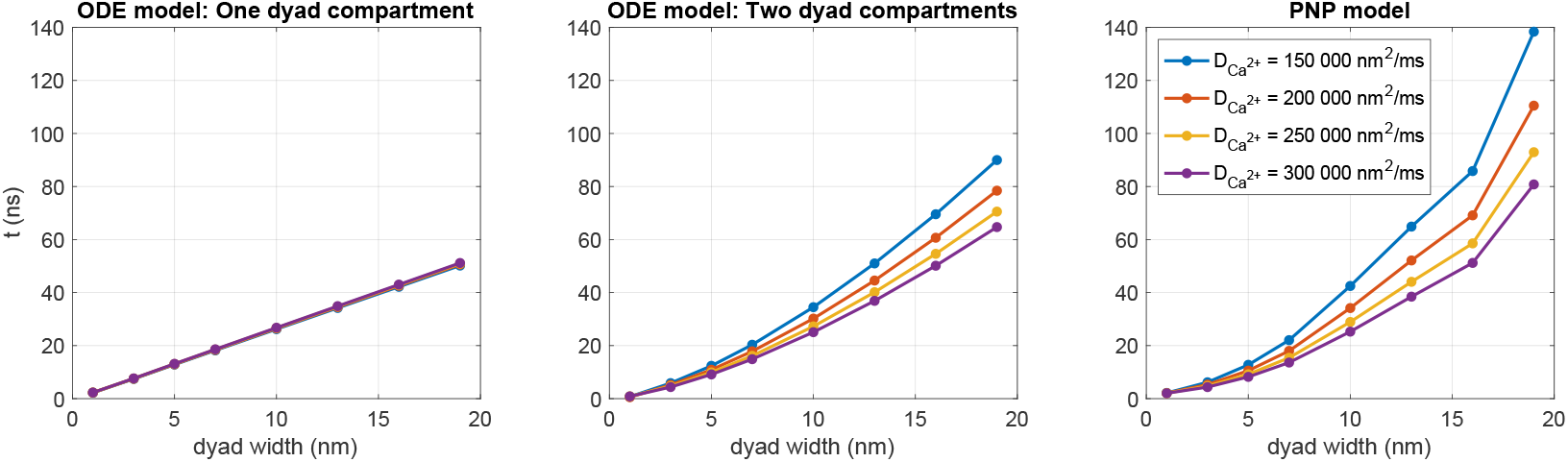
Time from the Ca^2+^ channel is opened until the [Ca^2+^] outside of an apposing RyR channel reaches 0.5 *µ*M in simulations of the two ODE models illustrated in Figure 18 and in the PNP model (left). We have used Δ*t* = 1 *µ*s. Note that in the leftmost panel, the results for different values of 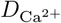 overlap

#### 3.11.2 ODE model representation with two dyad compartments

In an attempt to more realistically capture the RyR activation time in an ODE model representation, we extend the ODE model illustrated in Figure 18A to include two different dyad compartments, with associated concentrations (see Figure 18B). One of the dyad compartments, Ω_*c*_, is set up to represent the half of the dyad closest to the membrane (and the Ca^2+^ channel), and the other dyad compartment, Ω_*r*_, represents the half that is closest to the SR membrane (and the RyR). The two dyad compartments are connected by a diffusion flux, and the equations and parameters of the model are provided in the Supplementary Information.

The center panel of Figure 19, shows the RyR activation times for the ODE model with two dyad compartments. We observe that for this model, the RyR activation time increases as the diffusion coefficient is reduced, like also observed for the PNP model. In addition, the increase in the RyR activation time increases more rapidly as a function of the dyad width compared to the one dyad compartment case. However, the two compartment ODE model still underestimates the the increase in time as the dyad width is increased compared to the PNP model (right panel). The results of Figure 19 thus suggest that a PNP model or a spatially resolved reaction-diffusion model may be required to accurately capture the details of the Ca^2+^ dynamics in the dyad.

## 4 Discussion

### 4.1 Numerical solution of the PNP equations

Despite their ability to model essential biophysical processes, the PNP equations have received relatively little attention, primarily due to the significant computational cost associated with solving them. However, there is strong tradition for solving approximations of the system utilizing special geometries or symmetries or special features of the system, [44, 45], and the full equations have also been solved, see, e.g., [46, 23, 47, 48, 49, 24, 36]. In [24], for instance, we had to use a time step of Δ*t* = 0.02 ns to ensure numerical stability of the PNP system, making large-scale simulations impractical. In contrast, the numerical scheme employed in this study allows for a substantially larger time step of Δ*t* = 1000 ns, enabling simulations of physiologically realistic upstroke times of approximately 0.5 ms. The key to this 50,000-fold increase in time step is solving the entire system in an implicit and fully coupled manner, as shown in (31) and (32), rather than splitting the potential equation (1) and electrodiffusion equations (2). This change in numerical approach allows us to select the time step based on the required accuracy, rather than being constrained by the severe stability restrictions of the previous method. In fact, with our original scheme, the simulations presented here would have been computationally prohibitive, requiring infeasible computing efforts – defined here as exceeding a week per simulation. The computations reported here have been performed on a modest computer (a Dell Precision 3640 Tower with an Intel Core processor (i9-10900K, 3.7 GHz/5.4 GHz) with ten cores with two threads), and the computing time for, e.g., Figures 9–13 was 6 hours.

### 4.2 Electrodiffusion: What is the strongest term?

In Figure 5, we compare the magnitudes of the terms in the electrodiffusion equation (2) for a simple example of a perturbation from electroneutrality. The results indicate that electrodiffusion operates on two distinct time scales. Initially, the system rapidly establishes electroneutrality, during which the electrical term

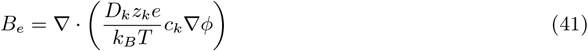

dominates the dynamics. This phase is characterized by a swift redistribution of ions driven by electrostatic forces, which act to neutralize local charge imbalances. Once electroneutrality is achieved, the system enters a slower relaxation phase, where the decay toward chemical equilibrium is governed by the diffusive term

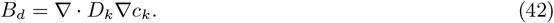

At this stage, the contribution of the electrical term *B*_*e*_ becomes negligible, and the remaining dynamics are primarily dictated by standard diffusion. From a global perspective, *B*_*e*_ is the strongest term when considering the entire simulation period. However, its dominance is transient; once electroneutrality is reached, the process becomes entirely diffusion-driven. This observation is in accordance with previous studies on electrodiffusion, see, e.g., [49], which emphasize the fundamental interplay between electrical forces and diffusion in neuronal environments and for other electrodiffusive processes.

### 4.3 Modeling ion channels

The PNP model is often implicitly used to represent the ionic fluxes through channels in the cell membrane in mathematical models. Ion channel fluxes cannot be represented by only taking diffusion into account; without the electrical force, the huge gradients in the ionic concentrations across the membrane would simply diffuse through ion channels and the cell would be useless. In most models, the current through an ion channel is modeled by assuming one-dimensional flow across the membrane and then integrating the Nernst equation analytically, often approximated by a model of the form (14). It is also possible to regard the channel as part of the computational domain and solve the PNP equations in the entire volume; the extracellular space, the cell membrane including the channel and the intracellular space. In Figure S1 and Figure S2 (Supplementary Information) we compare these approaches and find that the results are similar (as they should be), but not identical. The approach taken to represent the cross-membrane dynamics as part of the computational domain can be used for all the ion channels considered here, but not for the NCX. Modeling of the NCX has gained considerable attention and we have chosen to rely on the models in the literature. Since this flux does not follow directly from the PNP equations, we incorporate it using internal boundary conditions of the form (9) and (10). For consistency, we also treat the ion channel fluxes in the same manner.

### 4.4 Properties of electrodiffusion in the dyad

In Figures 10–13, we studied the ionic dynamics in the dyad when a Ca^2+^ channel, an NCX, or both were open in the dyadic cell membrane. In Figures 10 and 11, we observed that when the Ca^2+^ channel was open, the Ca^2+^ concentration increased considerably close to the Ca^2+^ channel and this also resulted in local changes in the other ionic concentrations to maintain electroneutrality. Moreover, the Ca^2+^ concentration increased across the dyad, potentially leading to the opening of RyRs on the membrane of the SR. Furthermore, we observed that the presence of an NCX in the dyad did not seem to significantly influence these dynamics. When the Ca^2+^ channel was closed, on the other hand, we observed that Ca^2+^ was transported into the dyad through the NCX. This resulted in a local increase in the dyadic Ca^2+^ concentration outside of the NCX (see Figure 12). Moreover, if we reduced the intracellular diffusion coefficient and the dyad width and placed the Na^+^ channel close to the NCX, the increased Ca^2+^ concentration across the dyad approached the threshold concentration of 0.5 *µ*M assumed to be required for RyR activation. Thus, under these conditions, an NCX could potentially trigger RyR opening in the absence of an open Ca^2+^ channel. A similar result was found for a reaction-diffusion model in [34].

In Figure 14, we further examined the properties of electrodiffusion in the dyad by considering the time from membrane Ca^2+^ channel opening until the concentration across the membrane (close to the RyRs) was high enough to trigger RyR opening. We observed that this time increased in a non-linear manner as the dyad width was increased. In addition, the time increased as the intracellular diffusion coefficient was decreased.

### 4.5 Simplifications in the representation of the dyad

In this study, we have applied the PNP equations to model ionic dynamics in the dyad. The PNP model has the potential of providing a more detailed and accurate representation of the electrodiffusion dynamics than the more commonly applied reaction-diffusion models. Our focus has been to use this detailed model for the dynamics in an otherwise relatively simple setup. Thus, simplifications have been applied to other aspects of the modeling representation. For example, we have only focused on the dynamics taking place before the activation of RyRs in the membrane of the SR. Therefore, the model used in this study does not include a representation of the SR or RyRs, but this would be a natural extension of the current framework. In addition, we have only included a single ion channel of each type and opened the ion channels deterministically at specific points in time. This approach could be extended to include several ion channels of each type and additional types of ion channels, membrane pumps and exchangers. Furthermore, a stochastic modeling approach could be applied for the opening of the ion channels, with an open probability depending on, e.g., the transmembrane potential. In this study, we have also only included two types of stationary Ca^2+^ binding buffers in the dyad, but other types of buffers, including mobile buffers could also be included (see, e.g., [28]). Additionally, we have applied a simple 3D rectangular geometric representation of the dyad instead of a detailed geometry and channel locations based on experimental imaging (see, e.g., [28, 50]). Another simplification in the present approach is that also the part of the domain that does not represent the dyad area (see, e.g., Figure 3) is represented by the same narrow intracellular space and the same Ca^2+^ binding proteins as those present in the dyad. A wider intracellular space and different Ca^2+^ binding proteins in this part of the domain would probably be more realistic.

### 4.6 Is it necessary to solve the full PNP system?

As shown in Figure 17, the reaction-diffusion model (39)–(40) provides a time-of-arrival estimate for the Ca^2+^-wave traveling from a calcium channel to the opposing RyR that is nearly indistinguishable from the result of the full PNP system (5)–(8), even though the two models (PNP and reaction-diffusion) exhibit some substantial differences close to the cell membrane (compare Figures 11 and 16). In particular, the reaction-diffusion model does not capture the altered concentration of other ionic species near the Ca^2+^ channel or the altered ionic concentrations (including the Ca^2+^ concentration) in the Debye layer. While it is theoretically possible to construct a scenario where a sustained electric potential gradient significantly influences ion transport even away from the cell membrane, such a setup would require carefully defined boundary conditions that may not be physiologically relevant. Based on our results, solving the reaction-diffusion equations appears sufficient for estimating time-of-arrival dynamics for the Ca^2+^ concentration in the dyad. In Figure 19, we observe that even a simple ODE model with two compartments can roughly capture the effect of a changed diffusion coefficient or dyad width on the time of arrival, but not to the same degree of accuracy as the reaction-diffusion model. Moreover, a single-compartment model is insufficient to reproduce the effect of a changed diffusion coefficient.

### 4.7 Continuous vs. discrete modeling of the dyad

As mentioned above, the PNP system serves as a continuous model for electrodiffusion in the dyad. However, the dyadic volume is extremely small, and ionic concentrations – particularly for Ca^2+^ – are very low; in fact, the number of Ca^2+^ ions is often below one at any given instant, [51]. The average volume of the dyad has been measured to be 4.39 × 10^5^ nm^3^, [52], which is approximately 4 × 10^7^ times smaller than the volume of an adult cardiomyocyte (16 pL), [53]. To accurately simulate this volume, we employ a spatial resolution at the nanometer scale and a temporal resolution at the nanosecond scale.

Mathematically, the PNP equations remain well-defined even at very low concentrations, but their physical interpretation becomes less straightforward. In [51], it is argued that continuous models should be interpreted as time-averages of inherently stochastic processes. An alternative approach is to use particle-based stochastic models, [54], which explicitly represent individual ion movements. Such stochastic models are frequently used to study Ca^2+^ dynamics in small, confined spaces where ion numbers are low and concentration-based descriptions may fail, [55, 56, 57]. Comparisons between stochastic and deterministic models generally indicate that for sufficiently small volumes, stochastic discrete models are more accurate than standard reaction-diffusion descriptions, particularly when dealing with nanometer-scale Ca^2+^ domains, [51, 58]. However, these comparisons are typically performed against reaction-diffusion equations or pure diffusion models, not against the PNP system itself. This distinction is critical because reaction-diffusion models, unlike PNP, do not account for the electrical forces governing ion movement in confined spaces. The presence of local charge imbalances and electric fields at the nanometer scale may significantly influence ion dynamics in ways that neither stochastic nor reaction-diffusion models capture.

Our approach is to analyze the PNP equations while acknowledging that they represent a continuous approximation of inherently discrete events. The results must therefore be interpreted with the caveat that local charge densities do not necessarily reflect instantaneous ion positions but rather their expected distributions over time.

## 5 Conclusion

In this study, we present a nano-scale computational model of electrodiffusion in the dyadic space using the Poisson-Nernst-Planck (PNP) equations. Our results confirm that resolving electrodiffusion dynamics at the nanometer scale provides critical insights into ionic transport mechanisms. We applied an improved numerical scheme for the PNP equations, which enables longer time steps and longer simulations, while maintaining numerical stability. This allowed us to investigate key factors influencing Ca^2+^ arrival at the ryanodine receptors, showing that both the diffusion coefficient and the dyad width significantly impact the time-of-arrival dynamics in a non-linear manner. Furthermore, we demonstrated that the sodium-calcium exchanger may trigger ryanodine receptor activation in the absence of an open Ca^2+^ channel if the dyad is narrow, the intracellular diffusion coefficient is reduced, and a Na^+^ channel is present in the dyad. Our findings also confirm the formation of a Debye layer near the cell membrane, which cannot be reproduced without including the electrical terms in the PNP equations. Finally, we highlighted that cross-species ionic interactions in the dyad are purely electrical, meaning that they do not manifest in pure reaction-diffusion models. These results underscore the necessity of nano-scale modeling for accurately capturing ionic dynamics in the dyad. The choice of model depends on the specific application: if representation of the Debye layer and cross-species ionic interactions are required, the full PNP system should be applied. However, if this is not a concern, a much simpler reaction-diffusion model could be an adequate approximation.

## Disclosure of writing assistance

During the preparation of this manuscript, the authors utilized the ChatGPT4o language model to enhance the language quality for contributions from non-native English speakers. Subsequent to this automated assistance, the authors rigorously reviewed and edited the manuscript to ensure its accuracy and integrity. The authors assume full responsibility for the content of the publication.

## Supplementary Information

### S1 Representing ion channels as membrane pores

Figures S1 and S2 compares the results of simulations of an open K^+^ channel when the channel is represented as selectively open for diffusion (Figure 2A and Figure S1) and when the channel is represented by internal boundary conditions (Figure 2B and Figure S2). Note that Figure S2 displays the same results as Figure 7, but in Figure S2 the scaling of the colormaps are adjusted to be the same as for Figure S1.

When the channel is represented as selectively open for diffusion (Figure 2A and Figure S1), the concentration of all the ionic species except for the one that is able to move through the channel is set to zero. Furthermore, the background charge density, *ρ*_0_, is set up such that *ρ* is initially zero everywhere (including in the channel). Thus, *ρ*_0_ in the channel is set up to directly counter the initial condition set up in the channel. In the simulation displayed in Figure S1, the initial concentration of K^+^ in the K^+^ channel is set up as a linear function of *x* between the intracellular and extracellular concentrations. But, as observed in [24], other initial concentration profiles could have been selected, changing the dynamics close to the channel somewhat.

Furthermore, in the simulation displayed in Figure S1, the diffusion coefficient for K^+^ was set to *d*_K+_ = 1.66 · 10^4^ nm^2^/ms in the K^+^ channel. This value was selected because it made the duration of the dynamics similar to the simulation in Figure S2, where the K^+^ channel conductance was set to a physiologically realistic value of *g*_K+_ = 5 pS.

Comparing Figures S1 and S2, we observe that the dynamics appear to be quite similar for the two different channel representations. For example, the intracellular potential approaches a value of about −80 mV in what appears to be the same timespan. Moreover, the K_^+^_ concentration changes locally near the channel and the other ionic species are also changed in these locations to counteract the deviation from electroneutrality. Furthermore, at rest, a Debye layer is present in both cases. On the other hand, a few differences between the two cases are also evident. For instance, the time evolution of the intracellular K^+^ concentration 3.5 nm outside of the K^+^ channel (lower panel) is a bit different in the two cases. The charge density, *ρ*, and the potential, *ϕ*, in the K^+^ channel is also different between the two cases.

**Figure S1:**
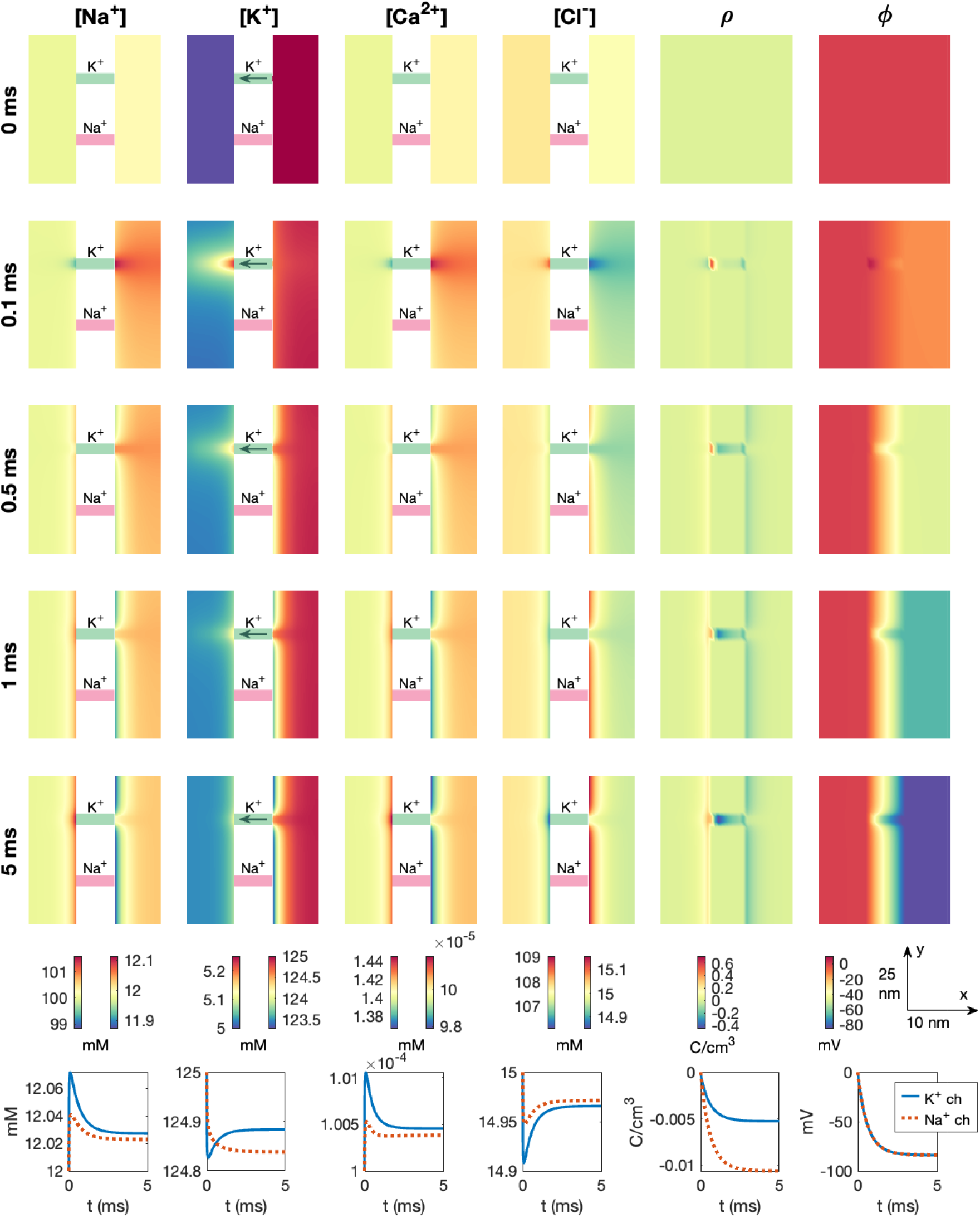
Dynamics following the opening of a K^+^ channel in a PNP model simulation with the channel represented like a membrane pore (see Figure 2A). We use Δ*t* = 1 *µ*s and an adaptive mesh like illustrated in Figure 3.

**Figure S2:**
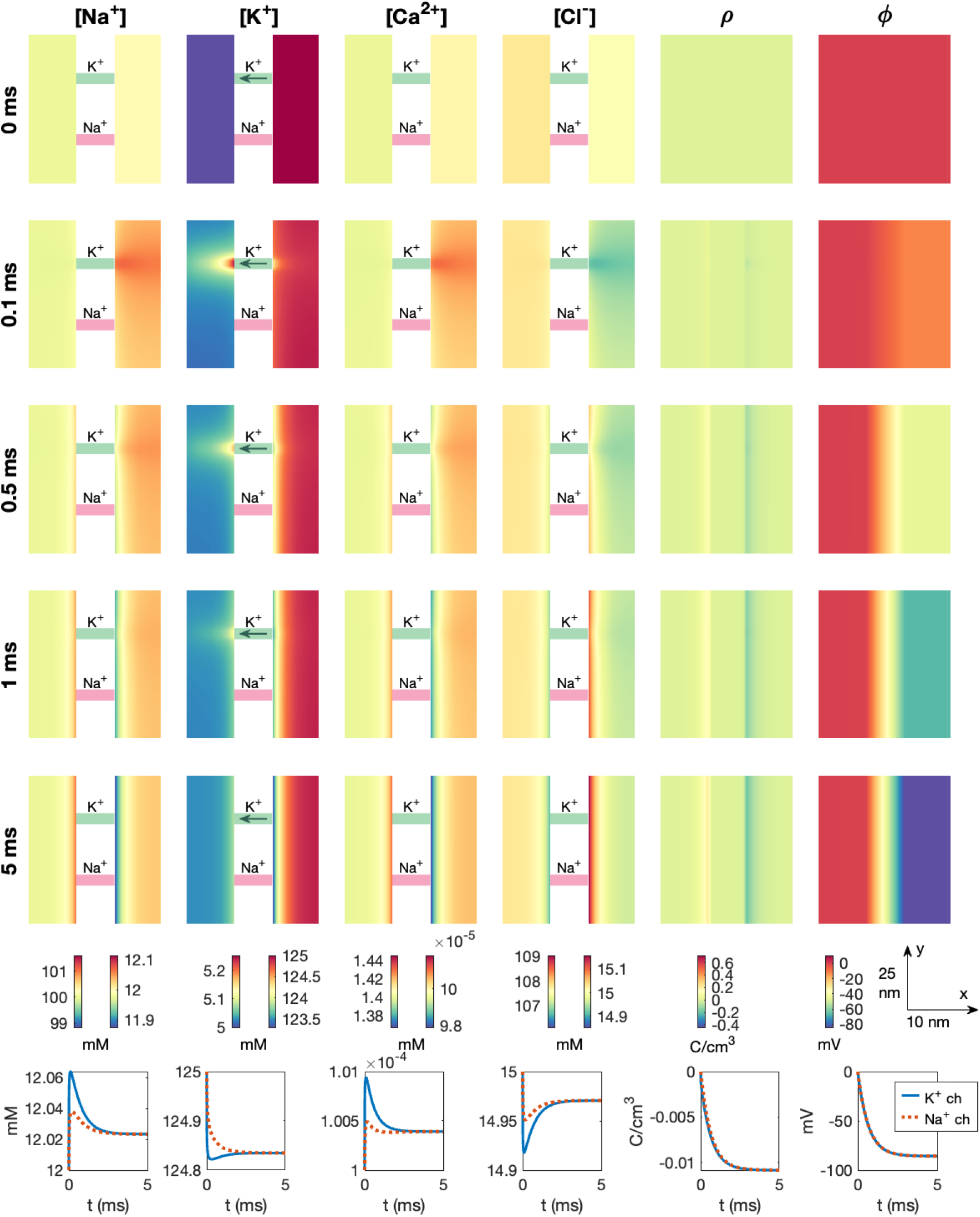
Dynamics following the opening of a K^+^ channel in a PNP model simulation with the channel represented using internal boundary conditions (see Figure 2B). This is the same simulation as in Figure 7, but the scaling of the colorbars are adjusted such that they match the ones used in Figure S1. We use Δ*t* = 1 *µ*s and an adaptive mesh like illustrated in Figure 3.

### S2 ODE model representations

In this supplementary section, the ODE model representations of the Ca^2+^ dynamics in the dyad that are compared to the PNP model are described.

#### S2.1 ODE model with one dyad compartment

The ODE model for one dyad compartment reads

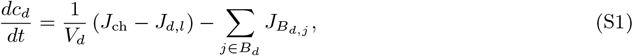

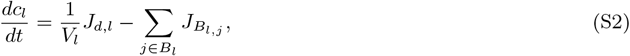

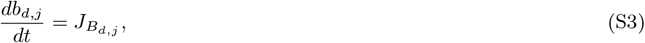

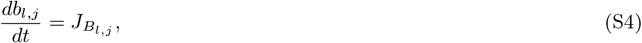

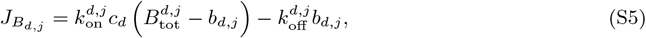

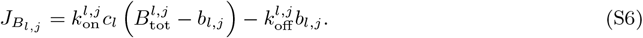

Here, *c*_*d*_ is the Ca^2+^ concentration and *b*_*d,j*_ are the concentrations of Ca^2+^ bound to a buffer in the dyad. Likewise, *c*_*l*_ is the Ca^2+^ concentration and *b*_*l,j*_ are the concentrations of Ca^2+^ bound to a buffer in the large cytosolic compartment. The concentrations have unit mM. The parameters *V*_*d*_ and *V*_*l*_ represent the volumes of the two compartments (in nm^3^), and *B*_*d*_ and *B*_*l*_ are collections of Ca^2+^ binding buffers in Ω_*d*_ and Ω_*l*_, respectively. To make the comparison to the PNP model more straightforward, we consider the same buffers as in the PNP model in both compartments (see Table 2).

The Ca^2+^ channel flux, *J*_ch_ (in mMnm^3^/ms), is defined as

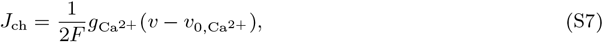

see (14). Here, 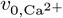 is the Nernst equilibrium potential of Ca^2+^ defined in (12). In the ODE model simulations, the intracellular concentration is defined as *c*_*d*_ and the extracellular concentration is fixed at the value provided for the extracellular Ca^2+^ concentration in Table 4. Furthermore, *v* is taken from the corresponding PNP model simulation.

The diffusion flux, *J*_*d,l*_ (in mMnm^3^/ms), is defined as

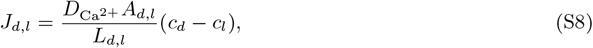

where 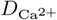 is the intracellular diffusion coefficient for Ca^2+^ (see Table 1), *A*_*d,l*_ (in nm^2^) is the average cross-sectional area connecting compartments *d* and *l*, and *L*_*d,l*_ (in nm) is the distance between the centers of the two compartments.

The parameter values of the ODE model are given in Table S1. Note that the dyad is assumed to be a volume of size

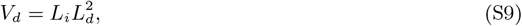

where *L*_*i*_ is the dyad width, and *L*_*d*_ defines the size of the dyad in the two other spatial directions. The value of *L*_*d*_ is fitted such that the time from Ca^2+^ channel opening to RyR threshold is roughly the same as for the PNP model for 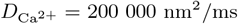 and *L*_*i*_ = 7 nm. Furthermore, *A*_*d,l*_ is defined as the intersection area between the dyad and the large compartment, i.e.,

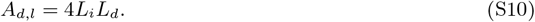

**Table S1:**
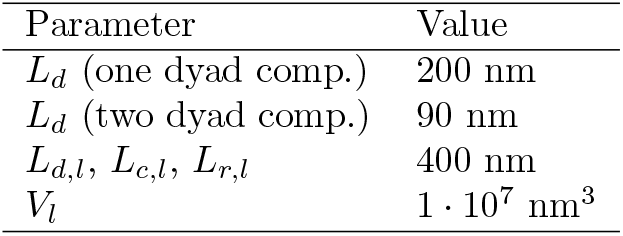
Parameter values used for the ODE model. The remaining parameter values are found in the tables of the main paper text. The value of *L*_*d*_ is fitted to match the RyR activation time of the PNP model in the case 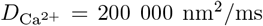 and *L*_*i*_ = 7 nm. The remaining parameter values are rough estimates.

| Parameter | Value |
| --- | --- |
| $L_d$ (one dyad comp.) | 200 nm |
| $L_d$ (two dyad comp.) | 90 nm |
| $L_{d,l}$, $L_{c,l}$, $L_{r,l}$ | 400 nm |
| $V_l$ | $1 \cdot 10^7$ nm<sup>3</sup> |

#### S2.2 ODE model with two dyad compartments

The ODE model for two dyad compartments reads

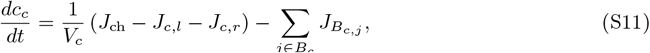

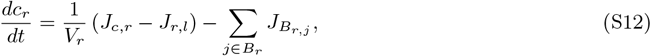

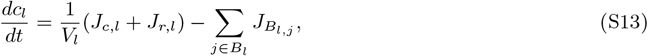

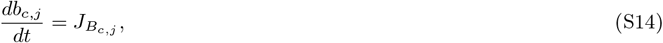

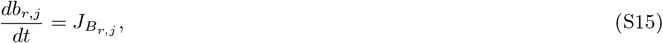

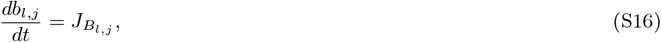

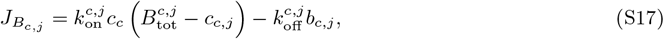

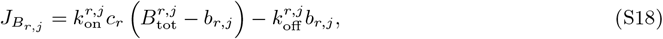

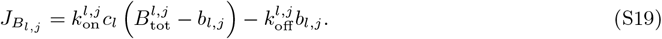

Here, *c*_*c*_ is the dyad Ca^2+^ concentration in the compartment close to the membrane Ca^2+^ channel, *c*_*r*_ is the dyad Ca^2+^ concentration in the compartment close to the RyR, and *c*_*l*_ is the Ca^2+^ concentration in the surrounding large cytosol compartment. The units and definitions of the remaining variables and parameters follow the same convention as for the ODE model with one dyad compartment.

The diffusion fluxes are defined as

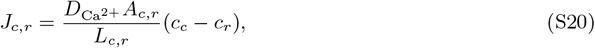

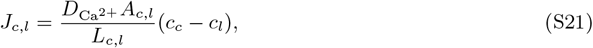

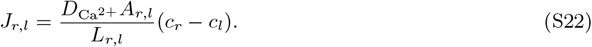

Moreover, we use the following assumptions regarding the compartment geometries,

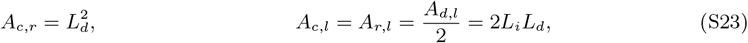

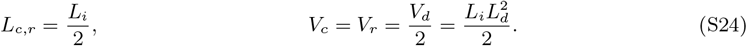

## References

[1] Mathis K Stokke, William E Louch, and Godfrey L Smith. Electrophysiological tolerance: a new concept for understanding the electrical stability of the heart. Europace, 26(11):euae282, 2024.

[2] Yoram Rudy and Jonathan R Silva. Computational biology in the study of cardiac ion channels and cell electrophysiology. Quarterly Reviews of Biophysics, 39(1):57–116, 2006.

[3] Yoram Rudy. From genes and molecules to organs and organisms: Heart. Comprehensive Biophysics, pages 268–327, 2012.

[4] Bogdan Amuzescu, Razvan Airini, Florin Bogdan Epureanu, Stefan A Mann, Thomas Knott, and Beatrice Mihaela Radu. Evolution of mathematical models of cardiomyocyte electrophysiology. Mathematical Biosciences, 334:108567, 2021.

[5] Thomas O’Hara, Laszl Virag, András Varró, and Yoram Rudy. Simulation of the undiseased human cardiac ventricular action potential: Model formulation and experimental validation. PLoS Computational Biology, 7(5):e1002061, 2011.

[6] Jun Chai, Johan Hake, Nan Wu, Mei Wen, Xing Cai, Glenn T Lines, Jing Yang, Huayou Su, Chun- yuan Zhang, and Xiangke Liao. Towards simulation of subcellular calcium dynamics at nanometre resolution. International Journal of High Performance Computing Applications, 29:51–63, 2015.

[7] Yuhui Cheng, Zeyun Yu, Masahiko Hoshijima, Michael J Holst, Andrew D McCulloch, J Andrew McCammon, and Anushka P Michailova. Numerical analysis of Ca2+ signaling in rat ventricular myocytes with realistic transverse-axial tubular geometry and inhibited sarcoplasmic reticulum. PLoS Computational Biology, 6(10):e1000972, 2010.

[8] Michael Nivala, Enno de Lange, Robert Rovetti, and Zhilin Qu. Computational modeling and numerical methods for spatiotemporal calcium cycling in ventricular myocytes. Frontiers in Physiology, 3(114), 2012.

[9] Piero C Franzone, Luca F Pavarino, and Simone Scacchi. Mathematical Cardiac Electrophysiology, volume 13. Springer, 2014.

[10] Natalia Trayanova and Gernot Plank. Bidomain model of defibrillation. Cardiac Bioelectric Therapy: Mechanisms and Practical Implications, pages 61–76, 2021.

[11] Natalia A Trayanova, Aurore Lyon, Julie Shade, and Jordi Heijman. Computational modeling of cardiac electrophysiology and arrhythmogenesis: toward clinical translation. Physiological Reviews, 104(3):1265–1333, 2024.

[12] Karoline Horgmo Jæger and Aslak Tveito. Efficient, cell-based simulations of cardiac electrophysiology; the Kirchhoff Network Model (KNM). NPJ Systems Biology and Applications, 9(1):25, 2023.

[13] Karoline Horgmo Jæger and Aslak Tveito. The simplified Kirchhoff network model (SKNM): a cell-based reaction–diffusion model of excitable tissue. Scientific Reports, 13(1):16434, 2023.

[14] Eugenio Ricci, Fazeelat Mazhar, Moreno Marzolla, Stefano Severi, and Chiara Bartolucci. Sinoatrial node heterogeneity and fibroblasts increase atrial driving capability in a two-dimensional human computational model. Frontiers in Physiology, 15:1408626, 2024.

[15] Karoline Horgmo Jæger, Verena Charwat, Kevin E Healy, Samuel Wall, and Aslak Tveito. Determining properties of human-induced pluripotent stem cell-derived cardiomyocytes using spatially resolved electromechanical metrics. The Journal of Physiology, 2025.

[16] Karoline Horgmo Jæger, James D Trotter, Xing Cai, Hermenegild Arevalo, and Aslak Tveito. Evaluating computational efforts and physiological resolution of mathematical models of cardiac tissue. Scientific Reports, 14(1):16954, 2024.

[17] Aslak Tveito, Karoline H Jæger, Miroslav Kuchta, Kent-Andre Mardal, and Marie E Rognes. A cell-based framework for numerical modeling of electrical conduction in cardiac tissue. Frontiers in Physics, page 48, 2017.

[18] Karoline H Jæger, Andrew G Edwards, Wayne R Giles, and Aslak Tveito. From millimeters to micrometers; re-introducing myocytes in models of cardiac electrophysiology. Frontiers in Physiology, 12:763584, 2021.

[19] Pietro Benedusi, Paola Ferrari, Marie E Rognes, and Stefano Serra-Capizzano. Modeling excitable cells with the EMI equations: spectral analysis and iterative solution strategy. Journal of Scientific Computing, 98(3):58, 2024.

[20] Zhilin Qu, Peter Hanna, Olujimi A Ajijola, Alan Garfinkel, and Kalyanam Shivkumar. Ultrastructure and cardiac impulse propagation: scaling up from microscopic to macroscopic conduction. The Journal of Physiology, 2024.

[21] Alfio Quarteroni, Toni Lassila, Simone Rossi, and Ricardo Ruiz-Baier. Integrated heart—coupling multiscale and multiphysics models for the simulation of the cardiac function. Computer Methods in Applied Mechanics and Engineering, 314:345–407, 2017.

[22] Sergio Alonso, Enrique Alvarez-Lacalle, Jean Bragard, and Blas Echebarria. Biophysical modeling of cardiac cells: From ion channels to tissue. Biophysica, 5(1):5, 2025.

[23] Jurgis Pods, Johannes Schönke, and Peter Bastian. Electrodiffusion models of neurons and extracellular space using the poisson-nernst-planck equations—numerical simulation of the intra-and extracellular potential for an axon model. Biophysical Journal, 105(1):242–254, 2013.

[24] Karoline Horgmo Jæger, Ena Ivanovic, Jan P Kucera, and Aslak Tveito. Nano-scale solution of the Poisson-Nernst-Planck (PNP) equations in a fraction of two neighboring cells reveals the magnitude of intercellular electrochemical waves. PLoS Computational Biology, 19(2):e1010895, 2023.

[25] Peter J. Mohr, David B. Newell, Barry N. Taylor, and E. Tiesinga. NIST reference on constants, units, and uncertainty. https://physics.nist.gov/cuu/Constants/index.html, Fundamental Constants Data Center of the NIST Physical Measurement Laboratory, 2018. Accessed: 2022-04-03.

[26] Maria G Kurnikova, Rob D Coalson, Peter Graf, and Abraham Nitzan. A lattice relaxation algorithm for three-dimensional Poisson-Nernst-Planck theory with application to ion transport through the gramicidin A channel. Biophysical Journal, 76(2):642–656, 1999.

[27] Andreas Solbrå, Aslak Wigdahl Bergersen, Jonas van den Brink, Anders Malthe-Sørenssen, Gaute T Einevoll, and Geir Halnes. A Kirchhoff-Nernst-Planck framework for modeling large scale extracellular electrodiffusion surrounding morphologically detailed neurons. PLoS Computational Biology, 14(10):e1006510, 2018.

[28] Johan Hake, Andrew G Edwards, Zeyun Yu, Peter M Kekenes-Huskey, Anushka P Michailova, J Andrew McCammon, Michael J Holst, Masahiko Hoshijima, and Andrew D McCulloch. Modelling cardiac calcium sparks in a three-dimensional reconstruction of a calcium release unit. Journal of Physiology, 590(18):4403–4422, 2012.

[29] Eleonora Grandi, Francesco S Pasqualini, and Donald M Bers. A novel computational model of the human ventricular action potential and Ca transient. Journal of Molecular and Cellular Cardiology, 48(1):112–121, 2010.

[30] Ada J Ellingsrud, Andreas Solbrå, Gaute T Einevoll, Geir Halnes, and Marie E Rognes. Finite element simulation of ionic electrodiffusion in cellular geometries. Frontiers in Neuroinformatics, 14:11, 2020.

[31] CN Fong and JAM Hinke. Intracellular Cl activity, Cl binding, and 36Cl efflux in rabbit papillary muscle. Canadian Journal of Physiology and Pharmacology, 59(5):479–484, 1981.

[32] James P Keener and James Sneyd. Mathematical Physiology. Springer, 2009.

[33] Mathieu Lemay, Enno de Lange, and Jan P Kucera. Effects of stochastic channel gating and distribution on the cardiac action potential. Journal of Theoretical Biology, 281(1):84–96, 2011.

[34] Glenn T Lines, Jørn B Sande, William E Louch, Halvor K Mørk, Per Grøttum, and Ole M Sejersted. Contribution of the Na+/Ca2+ exchanger to rapid Ca2+ release in cardiomyocytes. Biophysical Journal, 91(3):779–792, 2006.

[35] Karoline Horgmo Jæger, Verena Charwat, Bérénice Charrez, Henrik Finsberg, Mary M Maleckar, Samuel Wall, Kevin E Healy, and Aslak Tveito. Improved computational identification of drug response using optical measurements of human stem cell derived cardiomyocytes in microphysiological systems. Frontiers in Pharmacology, 10:1648, 2020.

[36] Karoline Horgmo Jæger and Aslak Tveito. Differential Equations for Studies in Computational Electrophysiology, volume 14 of Simula SpringerBriefs on Computing. Springer Nature, 2023.

[37] Mark B Cannell, CHT Kong, MS Imtiaz, and Derek R Laver. Control of sarcoplasmic reticulum Ca2+ release by stochastic RyR gating within a 3D model of the cardiac dyad and importance of induction decay for CICR termination. Biophysical Journal, 104(10):2149–2159, 2013.

[38] Dirk Gillespie. Recruiting RyRs to open in a Ca2+ release unit: Single-RyR gating properties make RyR group dynamics. Biophysical Journal, 118(1):232–242, 2020.

[39] Mark A Walker, George SB Williams, Tobias Kohl, Stephan E Lehnart, M Saleet Jafri, Joseph L Greenstein, W Jonathan Lederer, and Raimond L Winslow. Superresolution modeling of calcium release in the heart. Biophysical Journal, 107(12):3018–3029, 2014.

[40] Ivan Valent, Alexandra Zahradníková, Jana Pavelková, and Ivan Zahradník. Spatial and temporal Ca2+, Mg2+, and ATP2-dynamics in cardiac dyads during calcium release. Biochimica et Biophysica Acta (BBA)-Biomembranes, 1768(1):155–166, 2007.

[41] Mingwang Zhong and Alain Karma. Role of ryanodine receptor cooperativity in Ca2+-wave-mediated triggered activity in cardiomyocytes. The Journal of Physiology, 602(24):6745–6787, 2024.

[42] Thomas R Shannon, Fei Wang, José Puglisi, Christopher Weber, and Donald M Bers. A mathematical treatment of integrated Ca dynamics within the ventricular myocyte. Biophysical Journal, 87(5):3351–3371, 2004.

[43] G Faber and Y Rudy. Calsequestrin mutation and catecholaminergic polymorphic ventricular tachycardia: A simulation study of cellular mechanism. Cardiovascular Research, 75(1):79–88, July 2007.

[44] A Peskoff and DM Bers. Electrodiffusion of ions approaching the mouth of a conducting membrane channel. Biophysical Journal, 53(6):863–875, 1988.

[45] C Soeller and MB Cannell. Numerical simulation of local calcium movements during L-type calcium channel gating in the cardiac diad. Biophys J, 73(1):97–111, 1997.

[46] Tomasz Sokalski, Peter Lingenfelter, and Andrzej Lewenstam. Numerical solution of the coupled nernst-planck and poisson equations for liquid junction and ion selective membrane potentials. The Journal of Physical Chemistry B, 107(11):2443–2452, 2003.

[47] Jurgis Pods. A comparison of computational models for the extracellular potential of neurons. Journal of Integrative Neuroscience, 16(1):19–32, 2017.

[48] Allen Flavell, Michael Machen, Bob Eisenberg, Julienne Kabre, Chun Liu, and Xiaofan Li. A conservative finite difference scheme for Poisson–Nernst–Planck equations. Journal of Computational Electronics, 13:235–249, 2014.

[49] Jerzy J Jasielec. Electrodiffusion phenomena in neuroscience and the nernst–planck–poisson equations. Electrochem, 2(2):197–215, 2021.

[50] Thomas MD Sheard, Miriam E Hurley, John Colyer, Ed White, et al. Three-dimensional and chemical mapping of intracellular signaling nanodomains in health and disease with enhanced expansion microscopy. ACS nano, 13(2):2143–2157, 2019.

[51] Johan Hake and Glenn T Lines. Stochastic binding of Ca2+ ions in the dyadic cleft; continuous versus random walk description of diffusion. Biophysical Journal, 94(11):4184–4201, 2008.

[52] Takeharu Hayashi, Maryann E Martone, Zeyun Yu, Andrea Thor, Masahiro Doi, Michael J Holst, Mark H Ellisman, and Masahiko Hoshijima. Three-dimensional electron microscopy reveals new details of membrane systems for ca2+ signaling in the heart. Journal of Cell Science, 122(7):1005–1013, 2009.

[53] Anders Nygren, Céline Fiset, Ludwik Firek, John W Clark, Douglas S Lindblad, Robert B Clark, and Wayne R Giles. Mathematical model of an adult human atrial cell: the role of K+ currents in repolarization. Circulation Research, 82(1):63–81, 1998.

[54] Ulrich Dobramysl, Sten Rüdiger, and Radek Erban. Particle-based multiscale modeling of calcium puff dynamics. Multiscale Modeling & Simulation, 14(3):997–1016, 2016.

[55] Charin Modchang, Suhita Nadkarni, Thomas M Bartol, Wannapong Triampo, Terrence J Sejnowski, Herbert Levine, and Wouter-Jan Rappel. A comparison of deterministic and stochastic simulations of neuronal vesicle release models. Physical Biology, 7(2):026008, 2010.

[56] Claire Guerrier and David Holcman. The first 100 nm inside the pre-synaptic terminal where calcium diffusion triggers vesicular release. Frontiers in Synaptic Neuroscience, 10:23, 2018.

[57] María Hernandez Mesa, Kimberly McCabe, and Padmini Rangamani. Synaptic cleft geometry modulates nmdar opening probability by tuning neurotransmitter residence time. Biophysical Journal, 2025.

[58] GA Langer and A Peskoff. Calcium concentration and movement in the diadic cleft space of the cardiac ventricular cell. Biophysical Journal, 70(3):1169–1182, 1996.

